# The stem cell-type transcriptome of bioenergy sorghum reveals the spatial regulation of secondary cell wall networks

**DOI:** 10.1101/2023.04.22.537921

**Authors:** Jie Fu, Brian McKinley, Brandon James, William Chrisler, Lye Meng Markillie, Matthew J Gaffrey, Hugh D Mitchell, Galya Orr, Kankshita Swaminathan, John Mullet, Amy Marshall-Colon

## Abstract

Bioenergy sorghum is a low-input, drought-resilient, deep-rooting annual crop that has high biomass yield potential enabling the sustainable production of biofuels, biopower, and bioproducts. Bioenergy sorghum’s 4-5 m stems account for ∼80% of the harvested biomass. Stems accumulate high levels of sucrose that could be used to synthesize bioethanol and useful biopolymers if information about stem cell-type gene expression and regulation was available to enable engineering. To obtain this information, Laser Capture Microdissection (LCM) was used to isolate and collect transcriptome profiles from five major cell types that are present in stems of the sweet sorghum Wray. Transcriptome analysis identified genes with cell-type specific and cell-preferred expression patterns that reflect the distinct metabolic, transport, and regulatory functions of each cell type. Analysis of cell-type specific gene regulatory networks (GRNs) revealed that unique TF families contribute to distinct regulatory landscapes, where regulation is organized through various modes and identifiable network motifs. Cell-specific transcriptome data was combined with a stem developmental transcriptome dataset to identify the GRN that differentially activates the secondary cell wall (SCW) formation in stem xylem sclerenchyma and epidermal cells. The cell-type transcriptomic dataset provides a valuable source of information about the function of sorghum stem cell types and GRNs that will enable the engineering of bioenergy sorghum stems.

## Introduction

Optimized bioenergy crops are needed to provide feedstocks for a sustainable low carbon fuel economy. An ideal bioenergy crop, or ideotype, is one that can be produced sustainably, is high yielding, stress resilient, and compositionally optimized. Bioenergy sorghum (*Sorghum bicolor* L. Moench) is known for its drought and heat resilience and low input requirements, critical attributes for the production of bioenergy crops in marginal environments. In good environments, bioenergy sorghum hybrids have the genetic potential to accumulate ∼40-50 dry MT of biomass per hectare that can be converted to bioethanol or biopower with 75-90% greenhouse gas mitigation impact (Olson et al., 2012; Truong et al., 2017), an energy producing ratio (output/input) > 20 (Byrt et al., 2011), a carbon intensity (C.I.) for bioethanol production of ∼17 g CO_2_ e/MJ (Kent et al., 2020), and net carbon sequestration in Southern and Lower Midwestern U.S. locations (Gautam et al., 2020). The yield of bioenergy sorghum in water limited environments is lower (∼10-25 MT/ha) indicating there is a significant opportunity to improve the productivity of this crop in adverse environments (Truong et al., 2017). Sorghum stems account for ∼80% of the harvested biomass (Olson et al., 2012). Bioenergy sorghum stems at harvest are typically 4-5 m in length, and stem height is an early indicator for selecting bioenergy sorghum genotypes with increased biomass yield (Dos Santos et al., 2020). Sorghum stem growth is regulated in a complex way by development (McKinley et al., 2016), auxin transport (Multani et al., 2003), brassinosteroid signaling (Hilley et al., 2016; Hirano et al., 2017), an AGCVIII kinase (Oliver et al., 2021), gibberellin (Ordonio et al., 2014), and shade avoidance signaling (Yu et al., 2021). Phytomers generated by the shoot apical meristem/rib meristem initially produce leaves followed by the activation of the stem intercalary meristem at the upper end of the pulvinus that generates cells required for stem internode growth (Yu et al., 2022). Cells generated by the intercalary meristem subsequently elongate and then xylem sclerenchyma cells and epidermal cells accumulate secondary cell walls (SCWs) that help prevent stalk breakage (Kebrom et al., 2017; Yu et al., 2022). Transcriptional regulation of the sorghum stem SCW pathway has been investigated through analysis of stem tissues collected at various stages of sorghum development (Hennet et al., 2020).

Plant organs contain distinct cell types, each programmed to carry out specific functions that collectively influence tissue/organ/plant phenotypes (Giacomello, 2021). For example, leaf morphology and function are modulated by leaf growth (cells comprising intercalary meristems), gas exchange (guard cells), C4 photosynthesis (mesophyll and bundle sheath cells), and long-distance sugar transport via the phloem (sieve elements and companion cells). Thus, elucidating the developmental programs and regulatory modules of individual cells comprising complex tissues is needed to understand biochemical and physiological processes that influence tissue and organ functions, and traits that integrate functions across the entire plant (Deal & Henikoff, 2011; Yaschenko et al., 2022). Epigenomics, transcriptomics, proteomics/phosphoproteomics, and metabolomics are being used to quantify the functions of gene/protein regulatory networks in organs, tissues, and cells during development and in response to environmental perturbations. Advances in Laser Capture Microdissection (LCM), microfluidics, and cell-sorting technologies are enabling researchers to isolate specific cell types for transcriptome analysis and nuclei for chromatin accessibility profiling. The acquired information allows researchers to identify transcription factor binding sites that distinguish transcriptional programs of diverse cell types. Application of these techniques to cells of bioenergy sorghum plants will provide a fine-scale blueprint of gene and cell-specific function, and insight into cell-specific differentiation, cell-to-cell and inter-organ signaling, transport, and response to environmental cues.

Fluorescence-Activated Cell Sorting (FACS) (Birnbaum et al., 2005), Isolation of Nuclei Tagged Associated Cell Type (INTACT) (Deal & Henikoff, 2011), Fluorescence-Activated Nucleus Sorting (FANS) (Zhang et al., 2008), and immunopurification of ribosome-associated mRNA (Zanetti et al., 2005) all require the construction of transgenic lines and availability of cell-specific promoters (Teixeira & Pereira, 2010; Rogers et al., 2012). Single-cell mRNA sequencing (scRNA-Seq) is a high-throughput technique to achieve single-cell transcriptomes (Tang et al., 2009; Klein et al., 2015; Macosko et al., 2015). Although scRNA-Seq has been applied to a handful of plant species (Satterlee et al., 2020; Farmer et al., 2021; Liu et al., 2021; Y. Wang et al., 2021), it provides the “minimal information” on cell spatial organization (Giacomello, 2021). In addition, rigid plant cell walls constitute a major challenge for all these techniques since the prolonged time in protoplasting solutions may trigger unintended stress responses, and in turn produce noisy transcriptomes (Liu et al., 2022). LCM is an alternative technique that circumvents some of the aforementioned drawbacks in other techniques, and is uniquely suited for spatial analysis in species where transformation is difficult or low-throughput. It precisely isolates cells using a laser beam guided by microscopy and is based on cell anatomical positions and morphological differences rather than labeling strategies and protoplasting. It is particularly amenable to plant species because of their clear cellular histological layout restricted by plant cell walls. LCM protocols for distinct cell types in multiple plant species have been developed and continue to be optimized (Takahashi et al., 2010; Anjam et al., 2016; Blokhina et al., 2017; Hua & Hibberd, 2019; Pires et al., 2022) since its first use in mammalian studies (Emmert-Buck et al., 1996; Schüitze & Lahr, 1998).

LCM has been broadly applied in multiple plant species to isolate cell types from organs or tissues (Nakazono et al., 2003; Brooks III et al., 2009; Shiono et al., 2014; Shi et al., 2021). In sorghum, LCM was used to isolate mesophyll and bundle sheath cells of leaf (Covshoff et al., 2013; Döring et al., 2016), cells that are located in the shoot apical meristem (Paterson et al., 2020), and epidermal and cortical/stele cells from roots (Koegel et al., 2013; Sivaguru et al., 2013; Calabrese et al., 2016). Most of these studies focused on the expression of targeted genes, rather than the genome-wide characterization of gene expression profiles. Although the sorghum stem is an important organ for mechanical support and transportation of water, nutrients, and signaling molecules, it has not been a target for LCM analysis. Thus, a comprehensive transcriptomic atlas of different stem cell types in sorghum is not available.

To help close the gap in our understanding of sorghum stem biology, we used LCM to isolate five different stem cell types from sweet sorghum (cv. Wray) that were collected for RNA-Seq analysis. Our analysis identified differentially expressed genes in each cell type and their underlying regulatory networks. The gene regulatory network that modulates SCW formation in a cell specific manner during development was investigated by applying the LCM-derived data to a developmental transcriptome analysis of SCW formation in sorghum phytomers (Kebrom et al., 2017; Yu et al., 2021). This high-resolution systems analysis is useful for understanding functions of genes/pathways/networks expressed in individual cells, and regulatory mechanisms that contribute to integrated phenotypes that impact stem traits.

## Results

### Sorghum stem cell types have distinct transcriptomes

Cell types have distinct transcriptomes that are underrepresented and intermixed in whole tissue samples that combine cell types. To investigate sorghum (cv. Wray) stem cell-type specific transcriptomes, Laser Capture Microdissection (LCM) was used to collect five stem cell types (epidermis, pith parenchyma, phloem, vascular parenchyma, xylem sclerenchyma) from the middle of a fully elongated internode (phytomer 8) during the vegetative phase, 74 days after planting (DAP). RNA was isolated from each cell type for RNA-Seq analysis. UMAP analysis of the derived data showed that replicates (n = 4) of each cell type cluster together, and that different cell types form distinct clusters (Figure 1). In addition, the analysis showed that pith parenchyma and epidermal transcriptomes cluster near each other and far from cells derived from vascular bundles that form a separate group (Figure 1).

**Figure 1.**
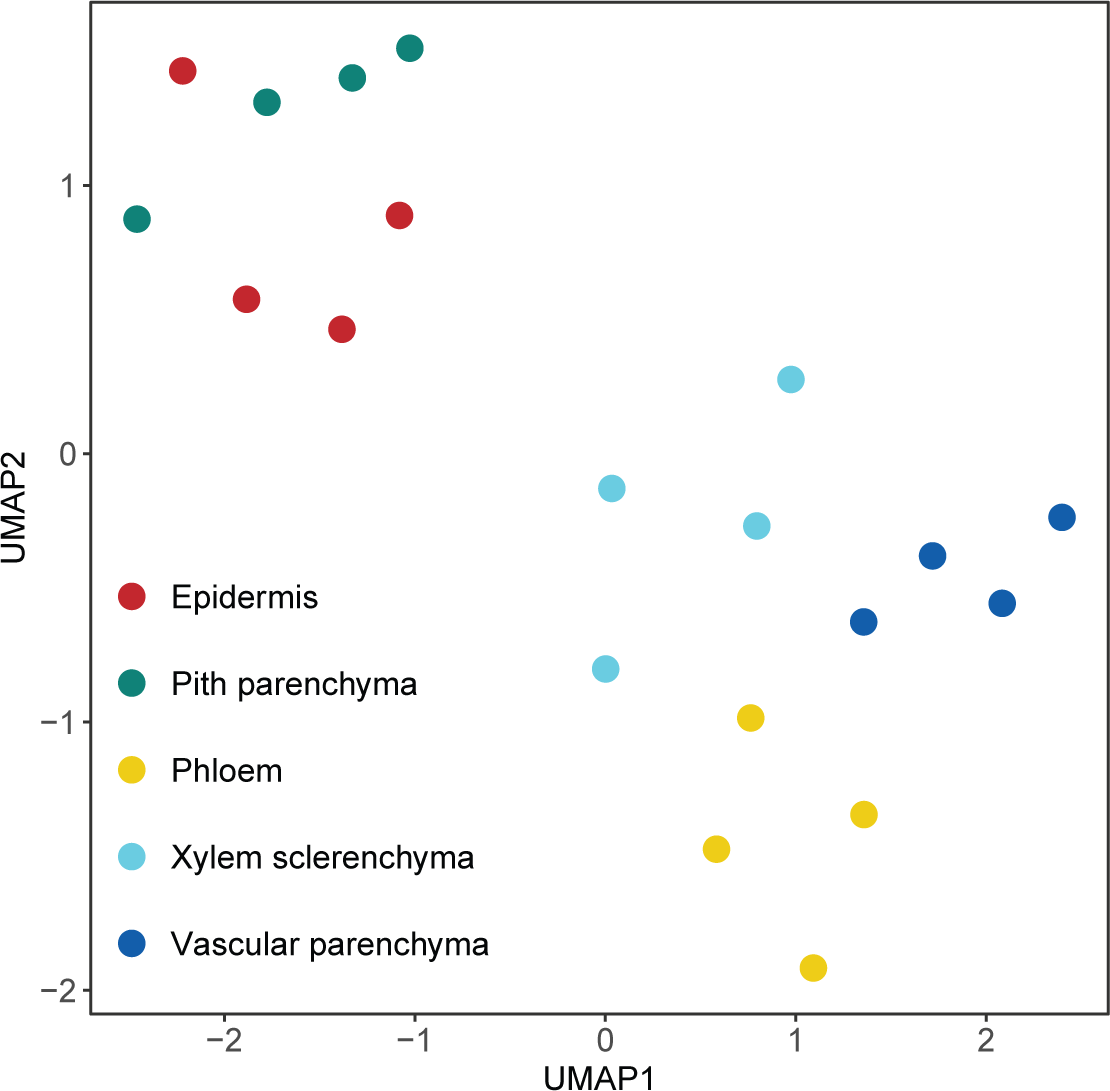
Dimension reduction UMAP plot showed a clear separation of five stem cell types, which form two large groups of vascular bundle cells (phloem, vascular parenchyma, and xylem sclerenchyma) and non-vascular bundle cells (epidermis and pith parenchyma).

While many genes are expressed in most cell types, genes with more selective expression in a single cell type are thought to contribute to cell-type specific functions and unique molecular characteristics. The Tau index (Yanai et al., 2005; Kryuchkova-Mostacci & Robinson-Rechavi, 2017) and Wilcoxon test were used to identify genes that were expressed exclusively (Tau = 1 & Wilcoxon p-value < 0.05) or predominantly (0.8 ≤ Tau < 1 & Wilcoxon p-value < 0.05) in each stem cell type. This analysis identified 708 genes expressed in a highly specific manner in the epidermis, 764 in the pith parenchyma, 562 in the phloem, 741 in the vascular parenchyma, and 327 in the xylem sclerenchyma (Figure 2 and Supplemental Data Set 1).

**Figure 2.**
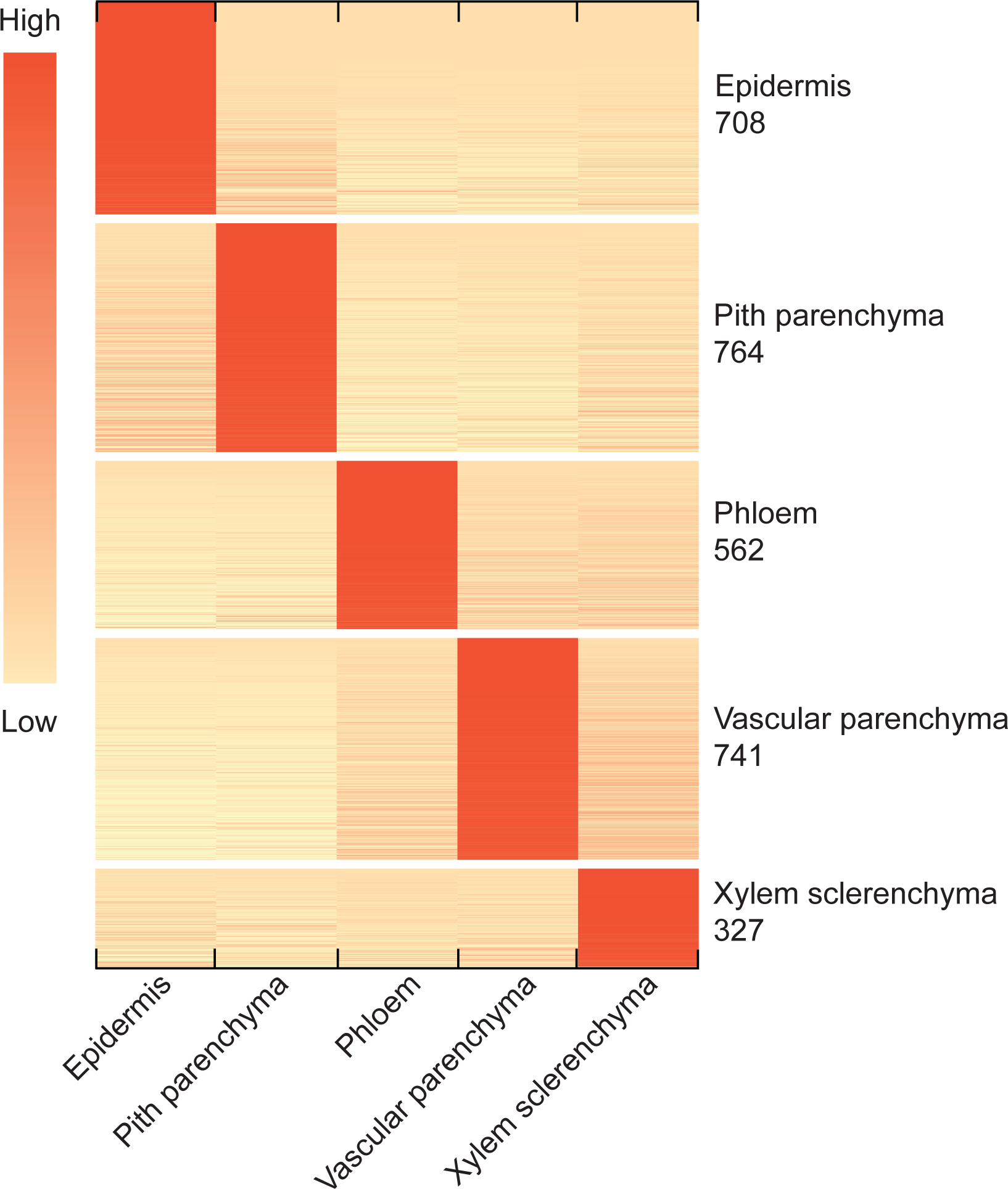
Cell-type specific genes were identified from the LCM-derived cell-type transcriptome dataset by applying the Tau index and Wilcoxon test. In the heatmap, gene expressions were centered and scaled by row.

The degree of specificity of the LCM-derived stem cell-type transcriptomes was investigated using known cell-type specific marker genes. Transcripts for numerous cell-type marker genes identified in other studies were present in the expected LCM-derived sorghum stem cell types (Supplemental Data Set 2). For example, the homolog of the *Arabidopsis thaliana LIPID TRANSFER PROTEIN 1* (*AT2G38540*, *LTP1*) that is expressed in epidermal cells (Thoma et al., 1994; Abe et al., 2001; Baroux et al., 2001) is expressed in LCM-derived sorghum stem epidermal cells (Supplemental Data Set 2). Similarly, *ALTERED PHLOEM DEVELOPMENT* (*AT1G79430*, *APL*) that is expressed in Arabidopsis inflorescence stem phloem cells (Abe et al., 2015; Schürholz et al., 2018) and *MYB DOMAIN PROTEIN 103* (*AT1G63910, AtMYB103*) and *SULFATE TRANSPORTER 1;2* (*AT1G78000*, *SULTR1;2*), genes that are expressed in xylem cells (Zhong et al., 2008), are differentially expressed in the corresponding LCM-derived sorghum stem cell types (Supplemental Data Set 2). No definitive marker gene has been reported for stem pith parenchyma cells (Shi et al., 2021); however, 66 genes with the pith-preferred expression pattern identified in a LCM study of the inflorescence stem in Arabidopsis (Shi et al., 2021) are expressed in a highly specific manner in sorghum stem pith parenchyma cells (Supplemental Data Set 2). These results indicate that the LCM-based approach used to enrich for specific stem cell types in the current study was effective. Several of the LCM ‘cell-type’ categories could be further subdivided using single-cell transcriptome analysis. For example, the LCM ‘epidermal cell’ category likely includes ground, cork, and possibly guard cells; the ‘phloem’ cell-type category includes sieve elements and companion cells; ‘vascular parenchyma’ cells likely include both phloem parenchyma and xylem parenchyma.

The utility of the LCM data was explored by examining gene expression differences in bulk stem tissue vs isolated cell types. Comparison of transcriptomes of a whole stem internode slice with LCM-derived cell types from an adjacent slice revealed that the majority of the cell-type specific genes identified have much lower expression in the whole tissue sample (Supplemental Figure 1), consistent with an expected dilution effect that occurs when cell-type RNA is collected and mixed during organ/tissue sampling. Thus, the cell-type specific RNA-Seq analysis increases the sensitivity of detecting transcripts derived from genes that are differentially expressed at low levels in specific cell types.

### LCM-derived cell-type transcriptomes uncover cell-type specific functions

#### Different functions distinguish vascular bundle and non-vascular bundle cells

The LCM-derived cell-type specific dataset was next used to investigate differences between cells that comprise vascular bundles (phloem, vascular parenchyma, and xylem sclerenchyma) and non-vascular bundle cells (pith parenchyma and epidermis) (Figure 1). Pairwise comparisons were made between all combinations of vascular and non-vascular cell types (Figure 3). An intersection of all up-regulated genes across vascular cell types compared to non-vascular cell types uncovered 127 common differentially expressed genes (DEGs) (FDR < 0.05 for calling DEGs in *edgeR*; Figure 3A and Supplemental Data Set 3). Gene Ontology (GO) enrichment analysis of these 127 genes revealed an overrepresentation of terms associated with localization and transmembrane transport (FDR < 0.05 for Fisher’s exact test; Figure 3A and Supplemental Data Set 4). The same analysis for down-regulated genes across vascular cell types (i.e. up-regulated genes across non-vascular cell types) uncovered 54 common genes (FDR < 0.05; Figure 3B and Supplemental Data Set 3), which are enriched for photosynthesis and light response GO terms (FDR < 0.05; Figure 3B and Supplemental Data Set 4). Thus, the up-regulation of transport processes distinguishes vascular cell types from non-vascular cell types, which are distinguished by the up-regulation of genes involved in photosynthesis.

**Figure 3.**
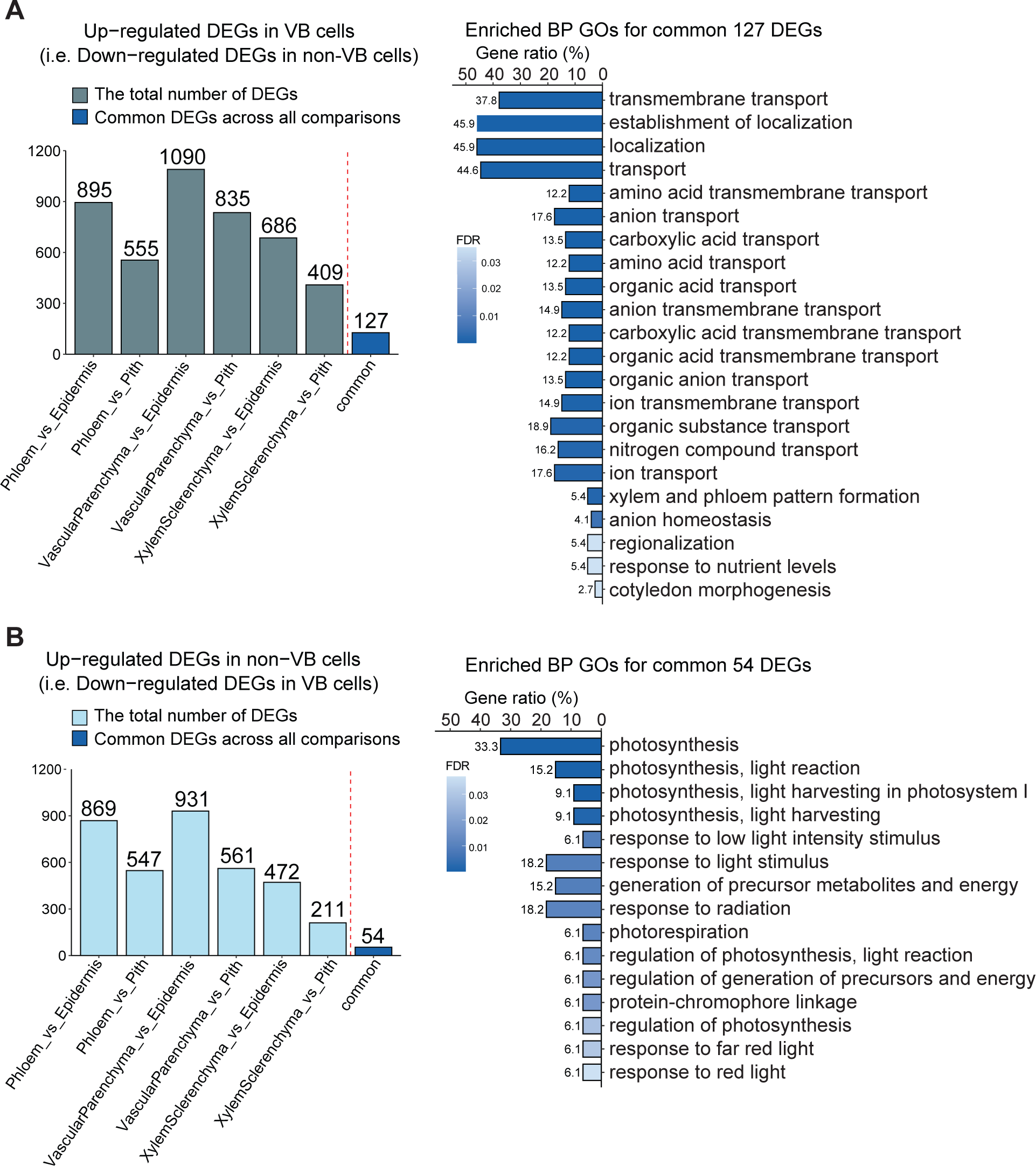
Common differentially expressed genes (DEGs) across cell-type pairwise comparisons of vascular bundle cells (phloem, vascular parenchyma, and xylem sclerenchyma) and non-vascular bundle cells (epidermis and pith parenchyma) imply that these two tissue types can be distinguished by transport-related and photosynthesis-related functions. (A) and (B) Up-regulated DEGs identified by *edgeR* (FDR < 0.05) and their enriched Gene Ontology (GO) Biological Process (BP) functions (FDR < 0.05) in vascular bundle (VB) cells (A) and non-VB cells (B). Bars labeled with ‘common’ are intersections of all pairwise comparisons.

Examination of differentially expressed transcription factors (TFs) revealed enrichment of different TF families in vascular and non-vascular cell-type gene regulatory networks (GRN) (Supplemental Data Set 13). GRN analysis found that the Basic Helix-Loop-Helix (bHLH) TF family has the most interactions with genes up-regulated in vascular cells, followed by the Homeodomain-leucine Zipper (HD-ZIP) family (Supplemental Figure 2A). In non-vascular cell types (epidermis and pith parenchyma GRNs), MYB TF family members are enriched (Supplemental Figure 2B). These results suggest that certain TF families play a role in controlling the functional differentiation between vascular and non-vascular cell types.

#### Different biological processes are up-regulated in each cell type

We further explored the transcriptomes of each of the five stem cell types by quantifying enriched GO terms of Biological Processes (BPs). GO enrichment analysis revealed common and unique biological processes across stem cell types (FDR < 0.05; Figure 4 and Supplemental Data Set 5). Pith parenchyma cells are significantly enriched for GO terms related to photosynthesis and light responses (Figure 4). Epidermal cells are enriched for processes involved in cell wall organization, fatty acid, lipid, and wax metabolism (Figure 4). Phloem and vascular parenchyma cells share processes for various transport and localization GO terms, but also have functions unique to each cell type: metal ion transport in phloem cells, and hormone-, terpenoid-, and glycosinolate-related processes in vascular parenchyma cells (Figure 4). Genes with differential expression in xylem sclerenchyma are enriched in GO terms for aromatic, phenylpropanoid, and lignin metabolic processes and share enriched GO terms with epidermal cells for cell wall organization and biogenesis (Figure 4).

**Figure 4.**
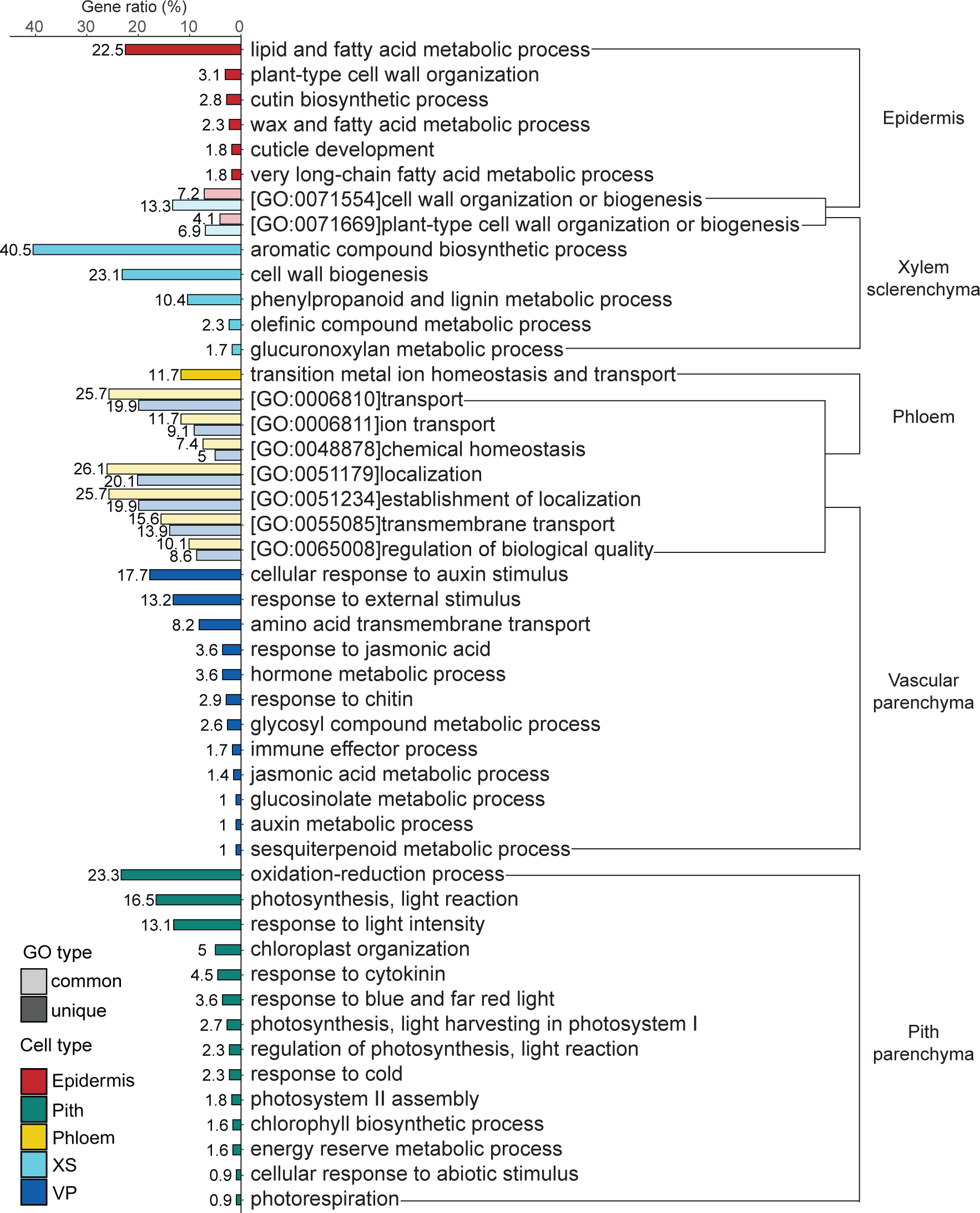
Cell-type specific genes are enriched in common (transparent colors) and unique (solid colors) Gene Ontology (GO) Biological Process (BP) functions (FDR < 0.05). Enriched GO terms were clustered if the overlap ratio of genes involved in these GO terms (Jaccard similarity coefficient) was more than 40%, followed by manually summarizing the function of each cluster (see method). Both GO clusters and orphan GO terms that cannot form a cluster are included in this plot. The X-axis represents the proportion of DEGs involved in this GO cluster or orphan GO term out of cell-type specific genes with GO annotations. The complete list of enriched GO terms can be found in Supplemental Table 5.

#### Different metabolic pathways and transporters are enriched in each cell type

Predicted enrichment of metabolic pathways in cell types based on the differential gene expression was examined using KEGG pathways (Kanehisa et al., 2021). The vascular cell types are enriched in secondary metabolic pathways and metabolism of terpenoids and polyketides (Figure 5 and Supplemental Data Set 6). Pith parenchyma specific genes are enriched for energy (photosynthesis) and carbohydrate metabolism (Figure 5 and Supplemental Data Set 6). Epidermal specific genes are enriched in metabolic pathways involving secondary metabolites and lipid, terpenoid, and polyketide metabolism (Figure 5 and Supplemental Data Set 6).

**Figure 5.**
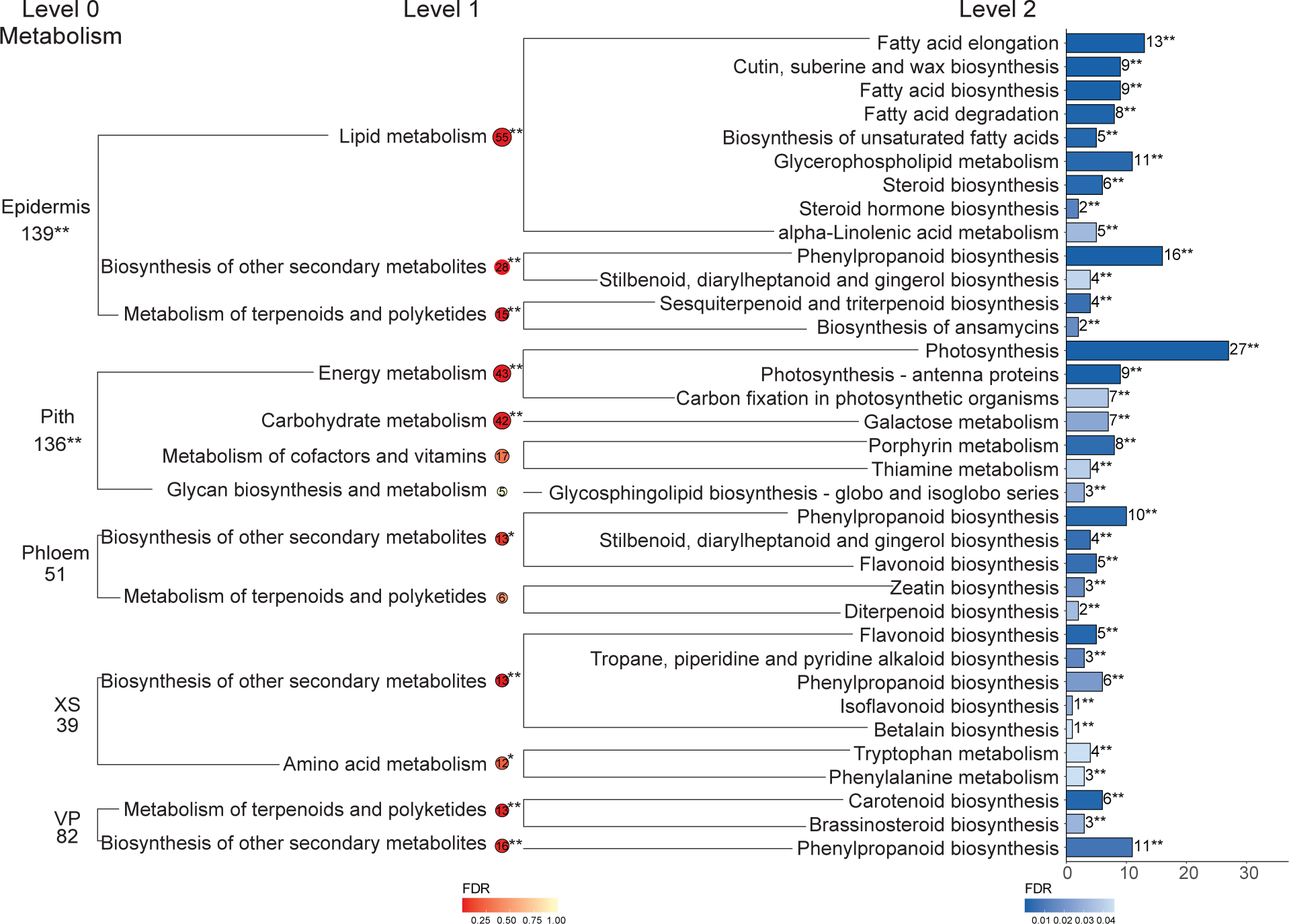
Cell-type specific genes are enriched in distinct metabolic processes defined in KEGG database. Pathways at different levels are shown as cell-type names for level0, bubbles for level1, and bars for level2, respectively. This plot only includes enriched level2 pathways (Fisher’s exact test, FDR < 0.05, labelled with “**”) due to the large number of defined pathways in KEGG, on which based its level1 pathway at the higher level is added into the plot or not. “**” and “*” represent significance levels of FDR < 0.05 and p-value < 0.05, respectively. Pathways (level1 and 2) and cell types (level0) without an asterisk indicate that they are not enriched. The complete result including gene identities and non-enriched pathways can be found in Supplemental Table 6. ‘VP’ represents ‘vascular parenchyma’ and ‘XS’ represents ‘xylem sclerenchyma’.

Genes with cell-type specific expression that encode transporters were also examined using KEGG annotations. Enriched transporter families mirror enriched metabolic pathways and biological processes across cell types (Figures 4-6 and Supplemental Data Set 5-7). The vascular phloem and vascular parenchyma cells have the largest number of cell-type specific genes encoding transporters, including electrochemical potential-driven transporters and sugar efflux transporters in both cell types (Figure 6 and Supplemental Data Set 7). The best match in sorghum for the sugar efflux transporters are SLC50A, SWEET; solute carrier family 50 in both cell types (Supplemental Data Set 7). The electrochemical potential-driven transporters include amino acid permeases, auxin efflux carrier family (PIN) proteins, and xenotropic and polytropic retrovirus receptor 1 (XPR1/PHO1) in both cell types (Supplemental Data Set 7). Genes encoding high and low affinity sulfate transporters (SULTR1 and SULTR2, respectively) and vacuolar iron transporter family proteins (VIT) are specifically expressed in vascular parenchyma cells, while a KUP system potassium uptake protein is exclusively expressed in phloem cells (Supplemental Data Set 7).

**Figure 6.**
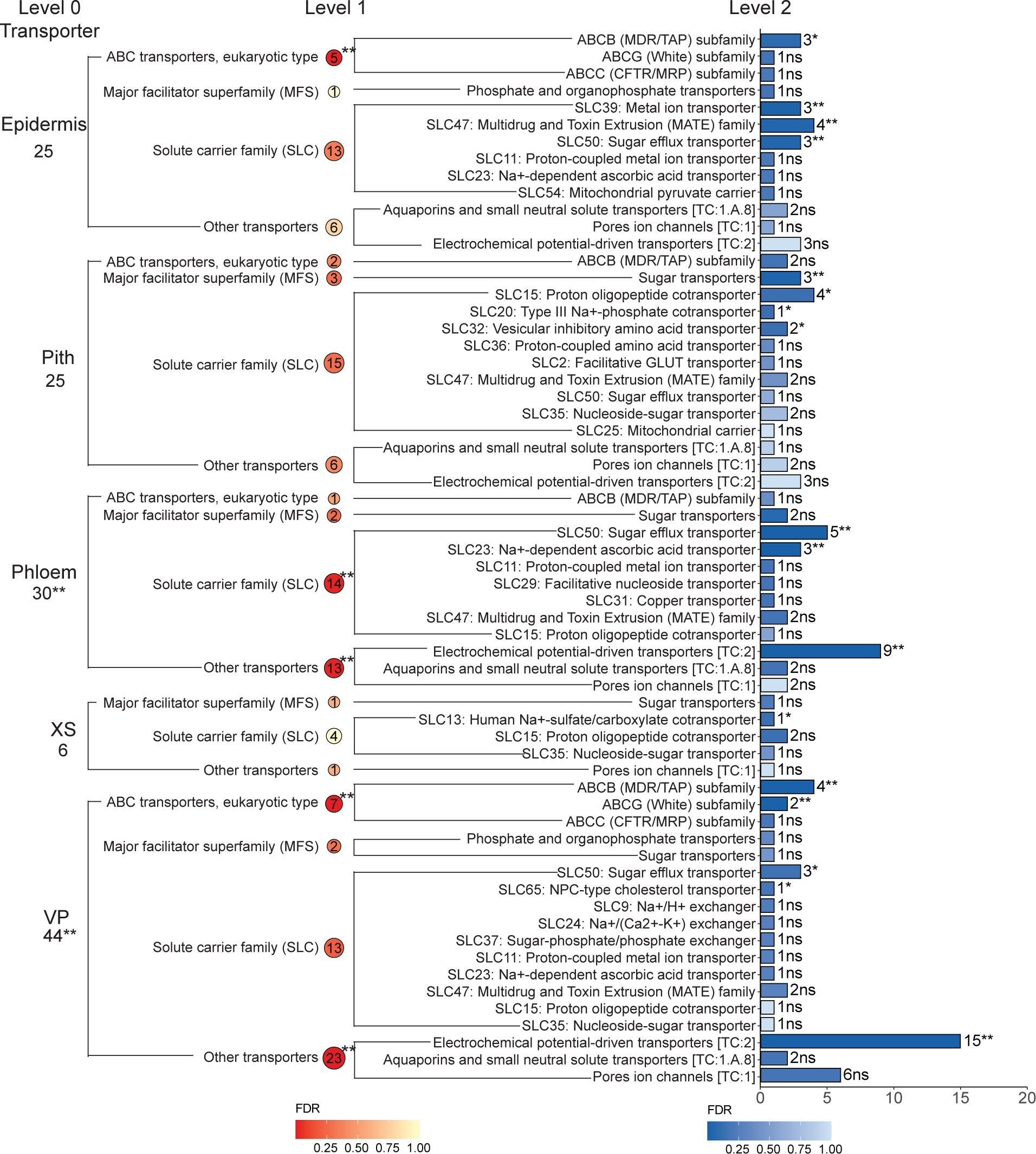
Cell-type specific genes are enriched in distinct transporter categories defined in KEGG database. Transporters at different levels are shown as cell-type names for level0, bubbles for level1, and bars for level2, respectively. This plot includes all the transporter categories regardless of enrichment significance, different from Figure5. “**”, “*” and “ns” represent significance levels of FDR < 0.05, p-value < 0.05, and not significant, respectively. The complete result including gene identities can be found in Supplemental Table 7. ‘VP’ represents ‘vascular parenchyma’ and ‘XS’ represents ‘xylem sclerenchyma’.

These results support prior knowledge about the flow of carbon and other cellular entities across cell types, but provide spatial resolution about specific processes that contribute to the distinction of different cell types within the sorghum stem.

### Cell-type <u>specific</u> gene promoters are enriched for unique TF binding motifs

Gene promoters can be bound by regulatory proteins from several different transcription factor families, and co-expressed genes are often commonly regulated by the same transcription factor(s) (Yin et al., 2021). Promoter sequences of cell-type specific genes identified in this study were analyzed to identify enriched cis-regulatory elements (CREs), including known and *de novo* motifs, within each cell type. Known motifs were identified using the *PlantPan 3.0* promoter analysis tool (Chow et al., 2019), and motif enrichment was determined using the Fisher’s exact test (FDR < 0.05); enriched *de novo* motifs were discovered using *MEME* (Bailey & Elkan, 1994) in *MEME* suite tools. Each cell-type specific gene set has significantly enriched motifs (Figure 7A). Nearly all epidermal specific genes (95.1%) have promoters that are enriched for the Basic-Leucine-Zipper (bZIP) TF family binding motifs, but are also uniquely enriched for the Zinc-Finger-Homeodomain (ZF-HD) and LBD binding motifs (Figure 7B). The majority (95.4%) of pith parenchyma specific gene promoters are enriched for C2H2 TF family binding motifs, but are also uniquely enriched in genes with MYB-related and TCP binding motifs in their promoters (Figure 7B). The GATA TF family binding motifs are overrepresented in vascular cells including phloem (89.9%), xylem sclerenchyma (87.8%), and vascular parenchyma (74.8%) (Figure 7B). The vascular cell types are also uniquely enriched in G2-like binding motifs (Figure 7B). These results indicate that distinct TF families may play a dominant role in regulating the cell-type specific expression of genes, and in turn control cell functions.

**Figure 7.**
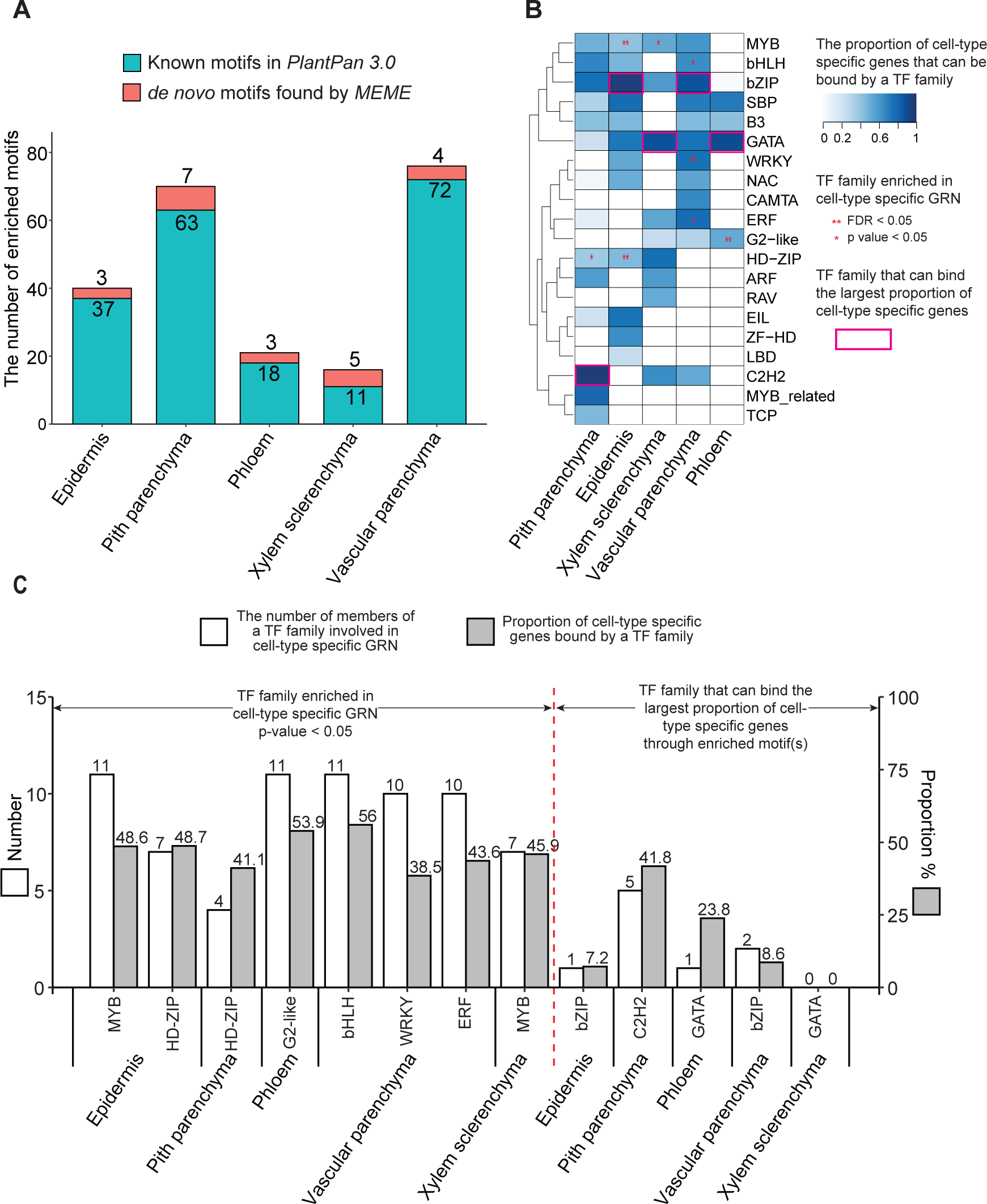
Transcription factor (TF) families could regulate cell-type specific genes by binding enriched motif(s) occurring in 1500bp promotor regions. (A) Cell-type specific genes are enriched in known motifs archived in *PlantPan 3.0* (green bars, Fisher’s exact test) and *de novo* motifs discovered by *MEME* (red bars). (B) Distinct TF families can regulate cell-type specific genes through binding known and *de novo* motif(s). Some TF families are enriched in cell-type specific GRNs (Fisher’s exact test). (C) TF families (labelled with asterisk in panel B) that are enriched in cell-type specific GRNs tend to directly regulate more cell-type specific genes than TF families (labelled with pink rectangle) that can bind the largest proportion of cell-type specific genes through enriched motifs.

### Cell-type specific TFs have direct and indirect modes of regulation

Promoter binding information was combined with gene co-expression analysis to construct cell-type specific GRNs to describe predicted TF-target gene relationships. Co-expression edges between nodes (genes) were estimated as Pearson Correlation Coefficients (PCC) (|PCC|< 0.8 & p-value < 0.05), and putative interactions between TFs and their targets were achieved using the *PlantPan 3.0* promoter analysis tool to identify CREs within the 1500bp promoter regions (upstream of start codon) of co-expressed genes. We first examined GRNs that only contain cell-type specific genes (0.8 ≤ Tau ≤ 1 & Wilcoxon p-value < 0.05; Supplemental Data Set 14), and found that TF families that are enriched in the GRNs (labelled with ‘*/**’ in Figure 7B) have regulatory edges with a large proportion (38.5%-56%) of the genes (Figure 7C, left panel). However, for the most part, these cell-type specific TFs do not belong to the TF families predicted to regulate the cell type specific genes based on the above promoter analysis (Figure 7B, framed with pink color). Thus, we expanded the GRNs to include non-cell-type specific TFs that are still highly expressed within the LCM samples to survey the full regulatory landscape of each cell type (Supplemental Data Set 15).

The expanded GRNs revealed that non-cell-type specific TFs, which are from the TF families with overrepresented CREs (Figure 7B), have regulatory edges with 64.5% to 97.4% of the cell-type specific genes, depending on the cell type (Figure 8, union of the Venn Diagram). Interestingly, these non-cell-type specific TFs are predicted to be regulated by cell-type specific TFs (Figure 8 and Supplemental Data Set 8), in which the promoters of the non-cell-type specific TFs have binding motifs belonging to cell-type specific TF families. These observations imply direct and indirect modes of regulation by cell-type specific TFs to turn on the spatially explicit expression of genes.

**Figure 8.**
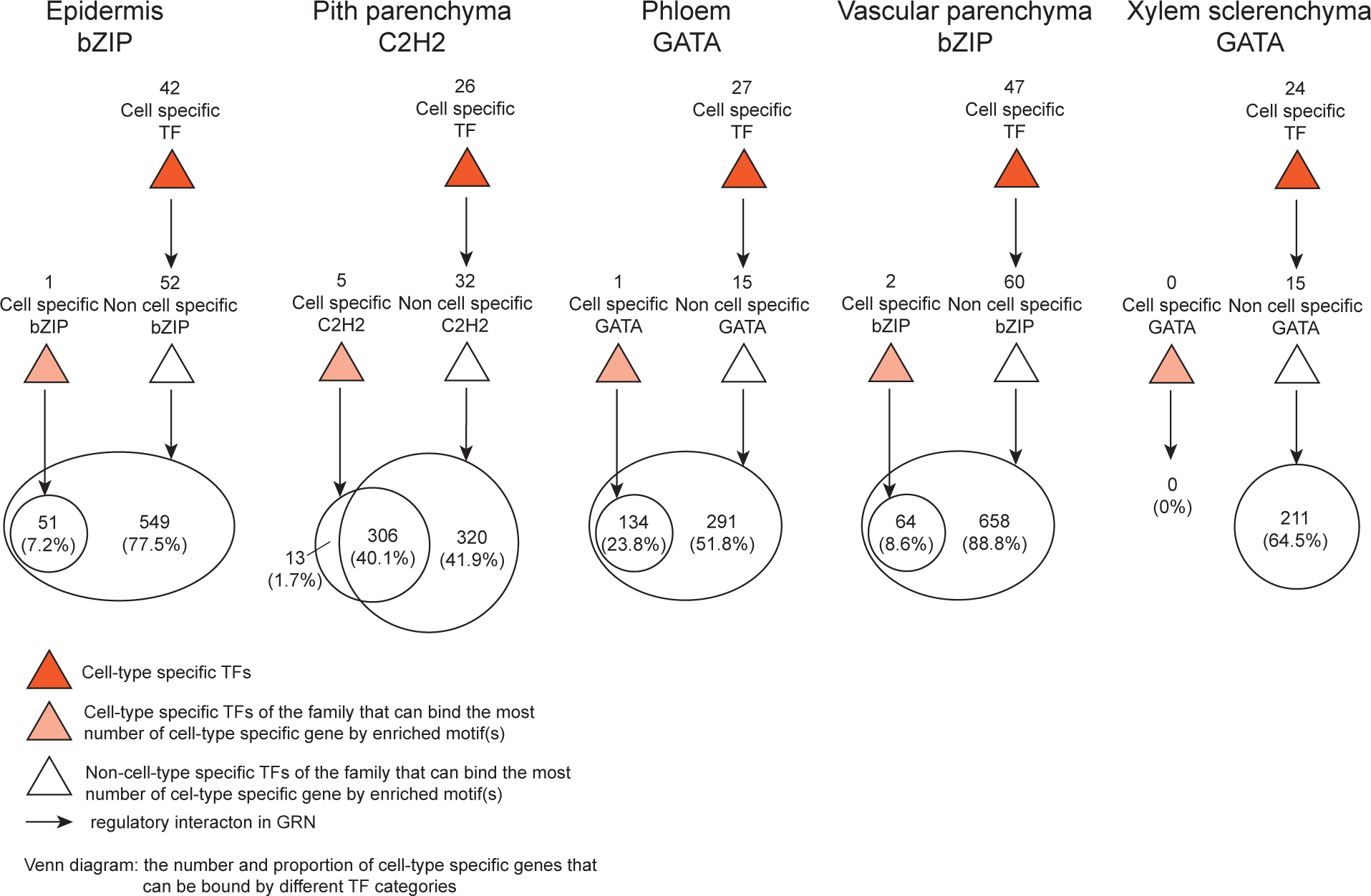
Cell-type specific TFs can regulate cell-type specific genes directly, or indirectly using non-cell-type specific TFs as intermedia. TF families involved in this plot are ones (labelled with pink rectangle in Figure 7B) that can bind the largest proportion of cell-type specific genes through enriched motifs.

### Stem secondary cell wall network analysis

Secondary cell walls (SCWs) are typically comprised of cellulose, hemicellulose, lignin, and cell wall proteins. Histological staining of lignin and cellulose in a cross section of the sorghum stem (Figure 9A) show that SCW formation is low in some stem cell types (i.e., pith parenchyma, phloem) and higher in other stem cell types (i.e., xylem sclerenchyma, epidermis). Moreover, SCW formation was previously shown to be repressed in the sorghum stem apical dome and intercalary meristems, regions of cell proliferation (Kebrom et al., 2017; Yu et al., 2021). SCW formation is activated concurrently with the onset of internode growth on cells that have completed elongation (Kebrom et al., 2017; Yu et al., 2021). To learn more about this complex pattern of SCW formation in the sorghum stem, we combined the spatial resolution of the LCM-derived stem cell-type specific transcriptome with the stem developmental profile of SCW formation (Kebrom et al., 2017; Yu et al., 2021).

**Figure 9.**
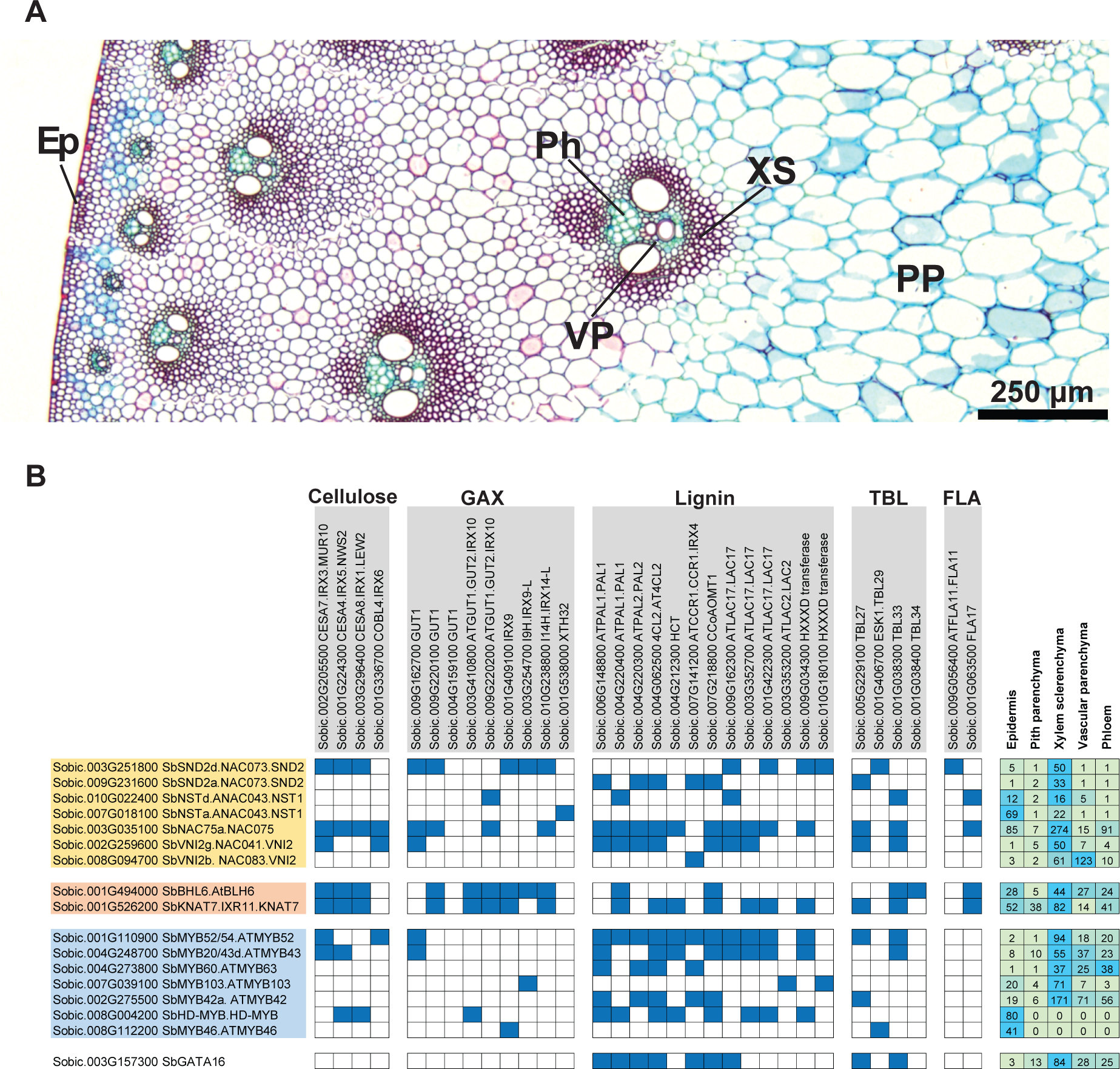
Secondary cell wall (SCW) TFs have cell-type preferred expression patterns and can differentially regulate genes involved in SCW formation, which helps explain the differential SCW pattern among cell types. **(A)** A FASGA-stained cross-section of the stem internode of Wray, the same as internodes collected for LCM-derived transcriptomes. Lignin was stained in red. ‘Ep’ for epidermis; ‘PP’ for pith parenchyma; ‘Ph’ for phloem; ‘VP’ for vascular parenchyma; ‘XS’ for xylem sclerenchyma. **(B)** The regulatory role and expression pattern of SCW TFs. Dark blue boxes represent regulatory interactions between SCW TFs (on the row) and SCW genes (on the column). SCW genes are grouped as biosynthetic pathways in which they are mainly involved. ‘GAX’ for glucuronoarabinoxylan; ‘TBL’ for Trichome Birefingence Like proteins; ‘FLA’ for Fasciclin-Like Arabinogalactan proteins. The heatmap on the right side is TMM-normalized TPM expressions of SCW TFs in the LCM-derived stem cell-type transcriptome dataset. Gene expressions are centered and scaled by row.

As the first step in the analysis, genes involved in SCW formation that are coordinately induced during stem development were identified using RNA-Seq data collected from nascent apical stem tissues and intercalary meristems where SCW formation is repressed, and stem tissue from elongating and recently fully elongated internodes where SCW formation is occurring (13 developmental time points) (Kebrom et al., 2017; Yu et al., 2021). *SbCESA4/7/8*, genes involved in SCW cellulose biosynthesis, were expressed at very low levels in apical, undeveloped stem tissues and in the growing zones of elongating internodes (i.e., internode 3, Int3-4, Int3-5) but at high levels in fully elongated stem tissues of internodes 3 and 4 (Int3-1, Int4) (Supplemental Data Set 9). Genes with developmental patterns of expression correlated with *SbCESA4/7/8* expression were identified using the same dataset (TPM>4 & PCC>0.91& FC>4). The 1082 genes that are co-expressed with *SbCESA4/7/8* included genes that encode enzymes involved in lignin and glucuronoarabinoxylan (GAX) biosynthesis, Trichome Birefingence Like (TBL) proteins, and FLA arabinogalactan-rich proteins that are known to contribute to SCW formation (Kumar et al., 2016; Coomey et al., 2020). In addition, numerous sorghum homologs of TFs involved in SCW formation were co-expressed with *SbCESA4/7/8* (i.e., *NST, SND, VNI2, MYB52*) (Hennet et al., 2020).

Genes involved in SCW formation were coordinately induced during stem development (Supplemental Data Set 10). Analysis of LCM cell-type transcriptome data showed that these genes are generally expressed at the highest levels in xylem sclerenchyma cells and at low levels in pith parenchyma cells, consistent with differences in accumulation of lignin, a marker for SCW formation, on the walls of these cell types (Figure 9A). Genes involved in SCW formation were also expressed at relatively high levels in the epidermis compared to pith parenchyma cells (Supplemental Data Set 10). GRN analysis was implemented to better understand how this complex pattern of cell-type specific SCW gene expression is regulated.

A GRN was constructed using the cohort of genes co-expressed with *SbCESA4/7/8* during stem development and used to identify predicted connections between TFs and genes involved in SCW formation (Supplemental Data Set 16). The connections between TFs and SCW genes are shown in Figure 9B, where predicted interactions are designated by blue boxes at the intersection between the TF and its downstream target SCW gene. This analysis showed that SbSND2d could potentially bind to the promoters of *SbCESA4/7/8*, several genes involved in GAX biosynthesis (i.e., *SbGUT1, SbIRX9*), *SbLAC17* and genes encoding HXXXD-domain acyltransferases such as PMT transferase (Petrik et al., 2014) that modify lignin. In contrast, SbSND2a was predicted to interact only with promoters of genes involved in lignin biosynthesis (i.e., *SbPAL, Sb4CL2, SbCCR1, SbCCoAOMT1*). SbNST1a and SbNST1d were predicted to have more limited interactions with the SCW gene promoters. SbNAC75 and SbVIN2g were highly connected to genes involved in cellulose, GAX and lignin biosynthesis. As expected, MYB factors (i.e., SbMYB52/54, SbMYB20/43, SbMYB60) had predicted connections to numerous genes involved in SCW formation including nearly every gene involved in lignin biosynthesis. During the analysis several additional transcription factors such as *SbGATA16* were identified that had predicted connections to subsets of genes involved in SCW formation (i.e., lignin pathway in the case of SbGATA16).

While the genes involved in SCW formation and the TFs they are predicted to interact with are co-expressed during development, the TF genes involved in SCW formation showed a variety of cell-type specific expression patterns. A heat map of TF expression in different stem cell types is shown to the right of Figure 9B. Most of the TFs are differentially expressed in xylem sclerenchyma cells consistent with differential accumulation of SCWs on this cell type (i.e., *SbSND2d, SbSND2a, SbVNI2g*). However, *SbNST1a* and *SbNST1d* were differentially expressed in xylem sclerenchyma and epidermal cells whereas *SbNAC75* was more highly expressed in xylem sclerenchyma than in epidermal cells. Expression of *SbMYB52, SbMYB43, SbMYB60*, *SbMYB103* and *SbMYB42* was highest in xylem sclerenchyma cells but expression was also relatively high in vascular parenchyma and phloem cell types. In contrast, *SbHD-MYB* and *SbMYB46* were highly expressed only in epidermal cells. Interestingly the sorghum homologs of genes that repress SCW formation in Arabidopsis (*SbBHL6, SbKNAT7*) (Liu et al., 2014) were expressed at high levels in most stem cell types, although *SbBHL6* expression was relatively low in pith parenchyma cells. These TFs had predicted interactions with genes involved in cellulose, lignin TBL and FLA synthesis. Taken together, sorghum stem TFs predicted to regulate genes involved in SCW formation show a variety of expression patterns across the stem cell types analyzed, suggesting that spatial expression of TFs as well as differential binding of TF-modules specifies the expression of genes involved in SCW formation in a stem cell-type specific manner. Potential TF x TF connections were also investigated to better understand the regulatory dynamics of the SCW network (Figure 10 and Supplemental Data Set 11). The analysis showed that TFs involved in SCW formation have many predicted interconnections including feedback loops (Supplemental Data Set 17). When connections were shown using an interaction matrix (Figure 10), some TFs showed no predicted connections with other TFs (SbSND2d, SbNST1a) while others were highly connected (i.e., SbSND2a, SbNAC75a, MYB52/54, GATA16). This analysis provides new information about the regulatory landscape of SCW formation in sorghum stem cell types.

**Figure 10.**
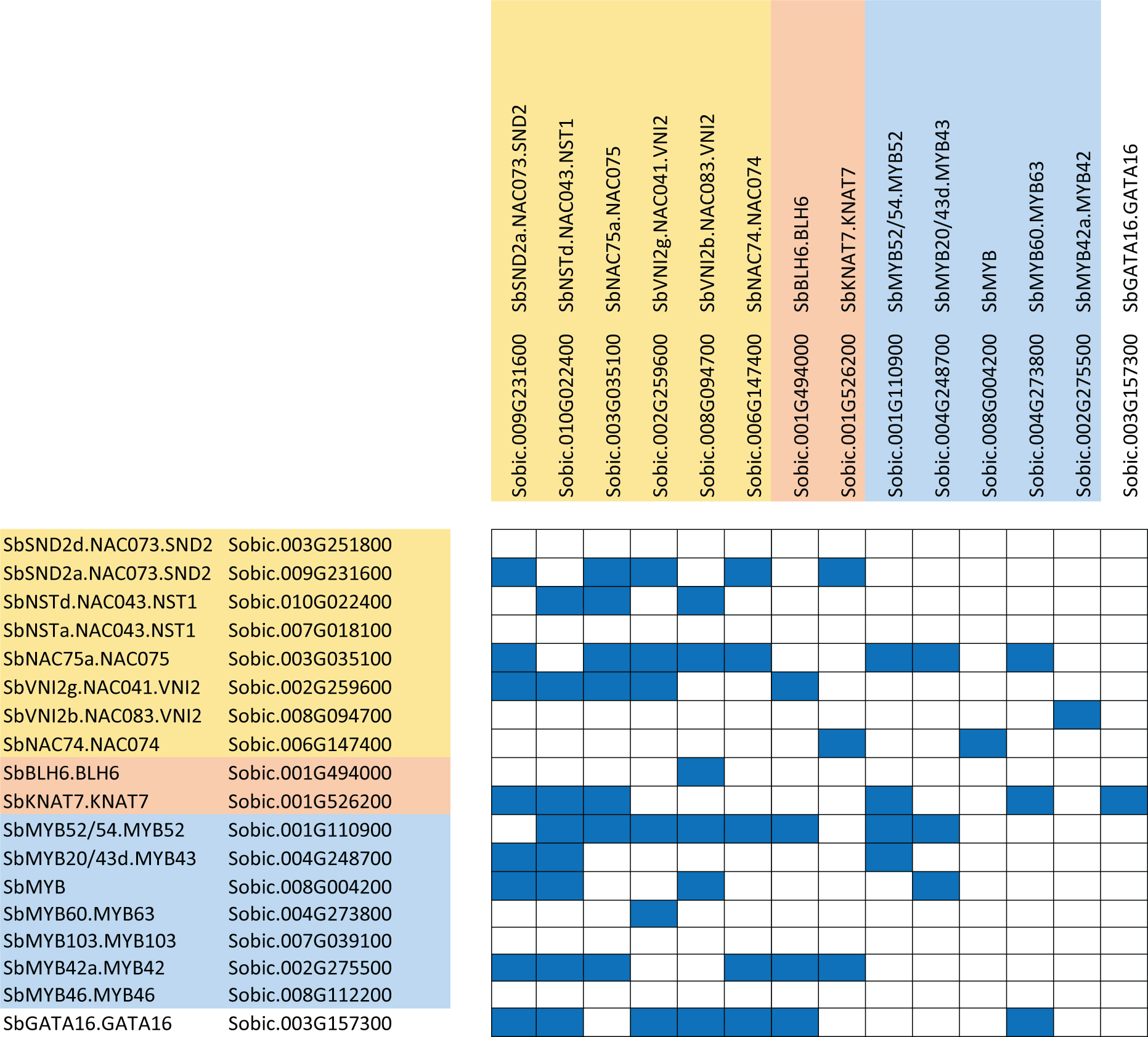
The regulatory landscape among SCW TFs. Dark blue boxes represent regulatory interactions among these TFs.

## Discussion

### A transcriptomic atlas of the sorghum stem

In this study, we dissected five cell types (epidermis, pith parenchyma, phloem, vascular parenchyma, xylem sclerenchyma) using LCM from the mid-internode of a sweet sorghum variety (cv. Wray) at the vegetative stage for genome-wide transcriptional analysis. The resulting high-resolution, spatial transcriptomes provide a valuable community resource that can facilitate research on sorghum stem biology. To our knowledge, this is the first comprehensive spatial analysis of grass stem transcriptomes. Single cell RNA-Seq (scRNA-Seq) was used in poplar stems to identify 20 distinct cell clusters (Chen et al., 2021), but scRNA-Seq has not been applied to sorghum. Similarly, in the model plant Arabidopsis, Shi et al. (2021) reported on high-resolution, spatial gene expression profiles of nine cell types in the inflorescence stem, investigated using Fluorescence-Activated Nucleus Sorting (FANS) and LCM. The analyses in Arabidopsis led to the isolation of more cell types due to the availability of species-specific cell markers that are not currently available for sorghum. However, the results of our analysis may enable the development of such marker lines for sorghum stem cell types. Likewise, the identification of highly cell-type specific genes (Tau = 1) will be useful for additional promoter element analysis to create molecular tools for cell-type specific expression of transgenes. Previous engineering attempts to increase sugar or lipid concentration in stem pith parenchyma cells of bioenergy grasses have not been highly successful (Wu & Birch, 2007; Watt et al., 2013; Jensen & Wilkerson, 2017), likely due to the lack of pith specific promoters (Wang et al., 2021).

### Spatial transcriptomes reveal distinct functions and metabolisms within cell types

Observed differences in spatial transcriptomes across the sorghum stem reflect distinct cell functions and unique molecular signatures of cells. By grouping transcriptomes into ‘vascular’ and ‘non-vascular’ cell types, we identified sets of genes that distinguish these two groups of spatially separated cell types based on biological functions. Vascular cell types (xylem sclerenchyma, vascular parenchyma, and phloem) commonly express genes involved in various transport processes, as expected, while non-vascular cell types (epidermis and pith parenchyma) commonly express genes involved in photosynthesis and response to light and radiation, which was not entirely expected. While stems of C4 grasses are often considered heterotrophic sinks (McCormick et al., 2009; McKinley et al., 2016), it has been known for decades that photosynthesis occurs in non-foliar tissues (Simkin et al., 2019). Stems may provide significant and alternative sources of photoassimilates essential for the synthesis of lipids required for growth, wax formation and optimization of yield (Hibberd & Quick, 2002), especially under stressful conditions such as drought (Ávila-Lovera et al., 2017). In fact, it was shown that tomato stems account for approximately 4% of whole plant photosynthetic activity (Hetherington et al., 1998). In *Brachypodium distachyon*, a model grass plant, the stem parenchyma cells (cortex parenchyma, cortical pith, and inner pith) have chloroplasts with thylakoid granum stacks. It was also shown that there is a reduction in plastid number from rind toward the stem center, and from younger to older stem internodes because with aging, chloroplasts in pith parenchyma cells are converted to amyloplasts (Jensen & Wilkerson, 2017). These observations are consistent with the differential expression of genes involved in photosynthesis in pith parenchyma cells in the vegetative sorghum stem in our study. Additionally, studies on stem photosynthesis have shown that carbon assimilation by Rubisco in stems is similar to the leaf but with CO_2_ diffusion through stomata in the stem, or by refixation of respiratory CO_2_ from the mitochondria (Simkin et al., 2019). While the specific genes that are upregulated in the epidermal and pith parenchyma cells suggest photosynthetic activity similar to leaves, we hypothesize that the overall contribution of the stem to atmospheric CO_2_ assimilation in sorghum is minimal, at least under non-stressful conditions. This hypothesis is supported by our observation that the expression of these photosynthesis genes across different tissue types in a related sorghum variety (McCormick et al., 2018; Arachchilage et al., 2020) are expressed at a much lower level (0%-8.1%) in stems compared to leaves pre-anthesis. However, specific studies aimed at quantifying stem photosynthesis are needed to learn more about the role of photosynthesis-related genes in the sorghum stem.

Besides photosynthesis, another enriched function for pith parenchyma cells is carbohydrate metabolism, which aligns with the role of pith parenchyma in carbohydrate storage post floral initiation in sweet sorghum stems. In sweet sorghum Della, the main forms of stem sugars before anthesis are monosaccharides such as glucose and fructose, but post-anthesis, monosaccharide sugar content decreases as sucrose and starch accumulate (McKinley et al., 2016). Similarly, our analysis in Wray shows that at the vegetative stage, starch and sucrose metabolism is not overrepresented in pith parenchyma cells (FDR < 0.05; Figure 5), but instead, galactose metabolism is the most overrepresented pathway (Figure 5). We found that pith parenchyma specific genes overlaid on the galactose pathway appear to direct molecular efflux into the production of D-Glucose, D-Fructose, and D-Galactose from precursor UDP-galactose by encoding unidirectional enzymes. Further analysis at additional developmental stages is needed to track changes in carbohydrate metabolism in Wray.

### The regulatory landscape contributes to the establishment of cell identity and function

The differentiation of specific cell types has been of great interest across biological domains. Cell type differentiation is attributed to a number of mechanisms from chromatin structure to peptide and hormonal signals (Pierre-Jerome et al., 2018). Of similar interest are the mechanisms that regulate cell-type specific gene expression in differentiated cell types that have different functions. Chromatin accessibility and DNA sequences (enhancers, promoters, and genic) have been shown to explain cell-type specific gene expression (Rosa et al., 2014; Uygun et al., 2019). Bioinformatics analyses have uncovered novel CREs in the promoters of cell-type specific genes (Uygun et al., 2017; Uygun et al., 2019) and used this information to identify transcription factors that bind to these elements and confer their cell-type specific expression (Noble et al., 2022). In this study, we used a bioinformatics approach that combined promoter CRE scanning and co-expression analysis to generate cell-type specific GRNs. As stated in the results and further discussed below, the vascular cell types are enriched in G2-like and GATA CREs, and the TF SbGATA16, a regulator of SCW formation, is highly expressed in xylem sclerenchyma cells (Figure 9B). In epidermal cells, seven TFs from the Homeodomain-Leucine Zipper (HD-ZIP) family are significantly overrepresented in the epidermis GRN (Figure 7B), and has predicted regulatory interactions with almost half of the epidermis specific genes (Figure 7C). Five of these seven epidermis-specific TFs are from the HD-ZIP class IV family (Javelle et al., 2011), of which most members have been shown to be preferentially expressed in the outmost protodermal/epidermis layer in multiple species (Lu et al., 1996; Abe et al., 2003; Javelle et al., 2011). Functional analysis of the 345 predicted target genes reveals that these epidermal specific HD-ZIP TFs regulate lipid and fatty acid biosynthetic pathways, which is the most statistically overrepresented biological function among epidermis specific genes (Figure 4 and 5). HD-ZIP family members are also enriched in the pith parenchyma specific GRN (Figure 7B), but none of these belong to the HD-ZIP class IV family (Javelle et al., 2011). This finding suggests that TFs from the HD-ZIP class IV family are specifically involved in epidermal expression of genes involved in lipid and fatty acid biosynthesis. Alternatively, the pith parenchyma expressed HD-ZIP TFs regulate nearly 40% of pith parenchyma expressed genes (Figure 7C), which are statistically enriched in photosynthesis-related biological processes. These results suggest that cell-type specific TFs, and even specific classes of TF families, play an important role in conferring unique expression profiles and functions of different cell types.

Through our bioinformatics analysis, we uncovered both direct and indirect interactions between cell-type specific TFs and cell-type specific genes, where the indirect interactions are mediated through non-cell-type specific TFs (Figure 8). Such a regulatory hierarchy has been identified in the past for Transforming Growth Factor Beta (TFG-β) in human cell lines (Mullen et al., 2011). It was found that the non-cell-type specific TFs Smad2 and Smad3 function in cell-type specific TFG-β signaling through the interaction with different cell-type specific TFs in embryonic stem cells, myotubes, and pro-B cells (Mullen et al., 2011). Such reaction chains can consist of various types of network motifs and have been shown to be prevalent in the regulatory landscape (Alon, 2007). In our GRN prediction, cell-type specific TFs and non-cell-type specific TFs can commonly regulate cell-type specific genes (Figure 8) by forming various network motifs (Supplemental Data Set 15). For example, Sobic.001G522500 (an epidermis specific bZIP) and Sobic.010G194900 (a non-epidermis specific bZIP) are predicted to mutually activate each other through the formation of a positive feedback loop that represses expression of Sobic.008G137500 (an epidermis specific gene). The Arabidopsis homolog of Sobic.008G137500 is AT2G37630, which is annotated to be involved in plant immune defense (Yang et al., 2008; Berardini et al., 2015). Studies have shown that immune defense is one of the main physiological functions for bZIP family members in plants (Jakoby et al., 2002). Also, homologs of these two sorghum bZIP TFs in Arabidopsis have been predicted to form a heterodimer (Deppmann et al., 2006). This possible heterodimer may act as a functional unit to regulate the plant defense associated gene. This is just one example of a hypothesized functional relationship between cell-type specific and non-cell-type specific TFs cooperatively regulating the expression of a cell-type specific gene, possibly in response to an environmental signal. Such relationships must exist to facilitate the complex but flexible regulatory network within distinct cell types.

### Regulation of genes involved in SCW formation in vegetative sorghum stems

The secondary cell wall biosynthetic pathways and their regulation have been studied at great depth in numerous plants (Taylor-Teeples et al., 2015; Kumar et al., 2016; Meents et al., 2018; Coomey et al., 2020). In bioenergy sorghum, SCW formation is of interest because biosynthesis of this specialized cell wall type in 4-5 m stems consumes a significant amount of photosynthate and contributes to stem strength and biomechanical properties that impact the propensity for stalk lodging (Gomez et al., 2017; Gomez et al., 2018). Genes involved in SCW biosynthesis and TFs that regulate SCW formation were previously identified in sorghum using phylogenetic analysis and tissue-level transcriptome profiling (Hennet et al., 2020). The current study extended the prior work by integrating transcriptome data derived from tissue level stem developmental analysis with the LCM analysis, and by using GRN analysis to predict targets of TFs that are involved in SCW formation. Sorghum stem developmental analysis identified 1082 genes that were co-expressed with *CESA4/7/8* genes that are involved in SCW formation. Thirty-two genes involved in cellulose, GAX and lignin biosynthesis and 17 genes encoding TFs (i.e., NST, SND, NAC075, BHL, KNAT7, MYB52) were co-expressed, consistent with their predicted involvement in SCW formation (Hennet et al., 2020). GRN analysis indicated that the TFs that regulate the expression of genes involved in CESA4/7/8, GAX and lignin are regulated in a complex combinatorial manner. The analysis also predicts sub-specialization of the two SND genes where SbSND2d was connected to genes involved in CESA, GAX and lignin biosynthesis but lacking connections to other TFs, whereas SbSND2a was connected to genes involved in lignin biosynthesis with several connections to other TFs in the network. NST1d was similarly connected to other TFs in the network, whereas NST1a was not. As expected, SbMYB60 was predicted to bind to genes involved in lignin biosynthesis (Scully et al., 2016). In contrast, SbMYB52 and SbMYB20 were predicted to bind to the promoters of genes involved in cellulose, GAX (GUT1) and lignin biosynthesis. The expression patterns of genes in the SCW network help to explain accumulation in the walls of xylem sclerenchyma and epidermal cells. Most genes in the network were more highly expressed in xylem sclerenchyma as expected. In addition, select SCW genes were also expressed in epidermal cells, or in two cases, differentially in epidermal cells compared to xylem sclerenchyma (i.e., *SbHD-MYB, SbMYB46*).

*SbBHL6* and *SbKNAT7* are part of the stem internode SCW developmental GRN and the corresponding TFs were predicted to have numerous connections to genes involved in cellulose, GAX and lignin biosynthesis. The Class II KNOX protein AtKNAT7 was found to repress secondary cell wall formation (Li et al., 2012) in part through interaction with BHL6 (Liu et al., 2014). KNAT7 also plays a regulatory role in conjunction with KNAT3 in xylem vessel formation (Wang et al., 2020). Expression of *SbKNAT7* was observed in all five stem cell types analyzed, with highest expression in xylem sclerenchyma and lowest expression in vascular parenchyma. *SbBHL6* was also expressed at high levels in all of the cell types except pith cells. It has been previously suggested that SbKNAT7 contributes to a negative feedback loop that functions to fine tune metabolic commitment to SCW formation, especially on interfascicular fibers (Li et al., 2012). SbKNAT7 and KNAT7:BHL6 heterodimers that can repress the commitment to SCW formation (Liu et al., 2014), could serve a similar function in sorghum stems. The repressing function of KNAT7 and BHL6 on SCW formation is further enhanced by interaction with OFP4, an OVATE FAMILY PROTEIN4 (Li et al., 2011; Liu & Douglas, 2015). It was therefore interesting to find that the sorghum homolog of *AtOFP4* (*SbOFP4*, Sobic.003G339100) was differentially expressed in stem pith parenchyma cells (Tau of 0.91; 6-40 fold higher expression than other cells), which accumulate only minimal levels of SCW. Selective and differential expression of *SbOFP4* in sorghum pith cells together with *SbKNAT7* and low levels of *SbBHL6* could help explain the repression of SCW accumulation on sorghum stem pith parenchyma cells relative to xylem sclerenchyma and epidermal cells.

## Methods

### Plant materials

*Sorghum bicolor* cv. Wray was grown in a greenhouse with the 14-h long day and well-watered conditions. Five-gallon pots were used for planting, filled with MetrtoMix900 (SunGro Horticulture, Agawam, MA) and fertilized with 30 g 14-14-14 Osmocote per pot (The Scotts Company, Marysville, OH). Seeds were obtained from the Texas A&M Breeding Program (College Station, TX, USA). Wray stem samples were collected from the most recently fully elongated internode (phytomer 8) of 74-day-old vegetative phase plants. Four biological replicates from four plants were collected, rapidly frozen, and sent on dry ice to Pacific Northwest National Laboratory (PNNL, WA, USA) for the isolation of different stem cell types using LCM, followed by RNA-Seq to achieve the cell-type transcriptomes (see below). Plant materials used for the stem developmental transcriptome dataset and the FASGA staining method for the stem cross-section to visualize the lignin component are described in Kebrom et al. (2017)

### LCM Protocol, RNA extraction, and RNA-Seq

Pre-cut and post-cut stem cell types by LCM visualized under microscope showed a high quality of target cell collections (Supplemental Figure 3), suggesting the efficiency of our developed LCM-isolation protocol. Internode sections of Wray were cut to 1 cm and flash-frozen in liquid nitrogen. The frozen sections were then embedded in optimal cutting temperature (OCT) compound and placed at −80°C for 30 minutes to fully solidify. The tissues were mounted by freezing onto magnetic chucks using UltraPure Distilled Water (Invitrogen 10977-015) and 20 μm sections were cut using a cryostat microtome with the following settings: Block temperature −20 °C and blade temperature −30 °C. Sections were placed on UV-treated membrane slides and the OCT was removed from the sections by dipping them in a serial dilution (70%, 85%, 100%) of ethanol at 4 °C for two minutes per dilution. Slides were then allowed to dry for 5 minutes and then either stored at −80 °C with desiccate or immediately placed on the LCM for sectioning using the following parameters: Energy 56 with delta 27, Focus 67 with delta −2, Cut Speed 20, and Magnification 20x (Blokhina et al., 2017). Working time at room temperature for each slide was no more than two hours. Five target cell types (epidermis, pith parenchyma, phloem, vascular parenchyma, and xylem sclerenchyma; Supplemental Figure 3A) after isolation were collected separately by catapulting into 0.2 mL tube caps containing 20 μL of RNAqueous Lysis Solution (Thermo AM1912), followed by total RNA isolation. Full-length cDNA synthesis, fragmentation of synthesized cDNA, and indexing were performed using SMART-Seq® v4 PLUS Kit (cat# R400753) for a RNA-Seq library construction, according to the manufacturer’s protocol. Single-end read sequencing with the read length of 150 bp was performed on NextSeq 550 Sequencing System using NextSeq 550 High Output v2 kit 150 cycles (cat#20024907). As a result, cell-type transcriptomes with the total read of 23870982 on average were generated. Bulk RNA-Seq for the whole stem tissue from the same internodes was performed following the same pipeline but without LCM-based target cell type enrichment process. Sequencing for the whole stem tissue generated the average library size of 31538064 reads.

### Read trimming and quality check for raw FASTQ sequencing files

*BBDuk* from *BBTools* suite (https://sourceforge.net/projects/bbmap/) was used to trim raw reads. *BBDuk* removed adapter sequences from the left end of reads, low-quality bases from both ends until the quality score at the operating base position reached the minimum of 10, and only kept reads with a self-defined minimal length and a quality score of 10 after the first two trimming steps. The minimal length in this study was defined as the one that reads equal to or longer than this length constitute 95% of total reads. Removing the remaining small part (5%), mainly consisting of extremely short reads, can guarantee that most of the sequencing information is retained but sequencing noise introduced by them is removed. Quality check was implemented using *FastQC* (Andrews, 2010) before and after the read trimming process to ensure that trimmed reads have no adapter content and reach the required quality score. Reads checked by *FastQC* after trimming were used for count quantification at the transcript level.

### Count quantification

*Salmon* (Patro et al., 2017) mapping-based mode was used to get transcript count. First, the decoy-aware transcriptome index was generated by concatenating the genome sequence to the end of the transcriptome sequence as described in its supportive alignment guide (https://combine-lab.github.io/alevin-tutorial/2019/selective-alignment/). Second, the *Salmon* quantifying method was used to get transcript counts. The transcriptome and genome sequence files used in the indexing process were from *Sorghum bicolor* v3.1.1 archived on *Phytozome* (https://data.jgi.doe.gov/refine-download/phytozome). 73.1%-83.2% of reads in cell-type transcriptomes after the read trimming were mapped onto the reference transcriptome by *Salmon*, depending on cell type. These transcript count profiles were used for downstream UMAP visualization, differentially expressed gene (DEG) analysis, Tau (τ) index calculation, and gene regulatory network (GRN) construction in cell-type analysis, but with different preprocessing and normalization processes (see below). Similarly, the average of 82.6% of reads in the whole stem RNA-Seq (n = 4) was mapped by *Salmon*. This transcript count for the whole stem was directly summed into gene count to compare with LCM-derived cell-type transcriptomes for the expression of cell-type specific genes.

### UMAP visualization

Uniform Manifold Approximation and Projection for Dimension Reduction (UMAP) was used to visualize LCM samples based on their expression profiles. (1) Transcript counts derived from *Salmon* were concatenated into gene counts. Genes were discarded if the number of samples, where the expression of a certain gene is detected (i.e. gene count > 0), was less than a ‘group size’. Group size is normally set as the number of replicates (i.e., n = 4 for this study).

However, the group size of 2 was used here to compensate for the possible adverse effect of the low sequencing depth on failing to capture lowly expressed genes. This adjustment leaded to a less stringent filtering criterion compared to the group size of 4. (2) Library size after gene filtering was scaled to one million. Gene counts were then normalized by log transformation. (3) A linear regression for each gene expression on the library size (before gene filtering) was constructed to remove the effect of library size on the gene expression. The residual of each linear regression was substituted for the log-normalized gene count achieved in (2) and used as the new proxy for the gene count. (4) Interquartile range (IQR) was used as the criterion to identify the top 2000 most variable genes. Only these genes were used for the downstream dimension reduction. (5) Principal Component Analysis (PCA) was performed to get the first 20 Principal Components (PCs), each of which was a multiple linear regression of these 2000 genes. (6) These 20 PCs were further reduced to two dimensions, on which the coordinates of all samples were visualized. The above steps were performed in *R* (version 4.0.5) using basic functions of ‘*lm*’ for linear regression, ‘*IQR*’ for interquartile range, and ‘*prcomp*’ for PCA, and using ‘*umap*’ library for UMAP.

### DEGs between vascular bundle cells and non-vascular bundle cells

*edgeR* (Robinson et al., 2010) was used to identify differentially expressed genes (DEGs) among pairwise comparisons between vascular bundle cells and non-vascular bundle cells. Transcript counts derived from *Salmon* were concatenated into gene counts as the input dataset for *edgeR*. A Gene filtering process was performed to remove low-expressed genes. The expressions of these genes were too low to be biologically significant or be translated into a functional protein. Specially, a gene was discarded if the number of samples, where the Count Per Million (CPM) of this gene is at least 5, was less than the group size of 2 (see UMAP visualization for group size definition). We normalized gene counts after the gene filtering process using the Trimmed Mean of M values (TMM) normalization method. DEGs were identified by performing the quasi-likelihood F-test (FDR < 0.05).

### Tau (τ) calculation and Wilcoxon test

Tau index (Yanai et al., 2005; Kryuchkova-Mostacci & Robinson-Rechavi, 2017) was used to assign cell-type specificity to each gene. Before calculating Tau, several steps were conducted to get normalized gene counts. These steps include (1) transcript counts from *Salmon* were used for the filtering process and TMM normalization, same with the processes in *edgeR* for identifying DEGs (see above), but at the transcript level; (2) transcript counts were first scaled by transcript length in the unit of kilobase, and then library size was scaled to one million; (3) transcript isoform counts were combined into gene counts, which will be used in Tau calculation. We performed gene concatenation after transcript count filtering and normalization because it can increase the accuracy by normalizing transcript count on its transcript length rather than gene length, particularly useful for some transcript isoforms with substantially different transcript lengths. The Tau calculating formula can be found in Kryuchkova-Mostacci et al. (2017). After getting Tau value for each gene, the Tau range of 0.8-1.0 was used to identify genes with cell-type preferred expression patterns, suggested by Kryuchkova-Mostacci et al. (2017). Since the Tau index does not account for variances among replicates, we applied the Wilcoxon test on each gene passing the Tau threshold to evaluate whether its expression was significantly higher in the cell type with the highest expression than that in other cell types (p < 0.05). Genes with the Tau of 0.8-1.0 and p < 0.05 in the Wilcoxon test were finally defined as ‘cell-type specific genes’ (Figure 2 and Supplemental Data Set 1). Among these, genes with the Tau of 1 were exclusively expressed in one cell type, thus having the highest specificity.

### BLAST to find the best hits in sorghum for known cell-type marker genes

Genes with known cell-preferred expression patterns were collected from literature across multiple species, including Arabidopsis, maize, and rice. *BLAST+* package (Camacho et al., 2009) was used to find the best sorghum hits for these known marker genes. Peptide sequences of these marker transcript isoforms in corresponding species and all sorghum transcript isoforms were retrieved using *Phytozome Biomart* tool (https://phytozome-next.jgi.doe.gov/biomart/martview/12933a5a373fa2cefc11af2ecd5dfba5). A sorghum protein database was constructed using *makeblastdb* function. Protein BLAST (i.e., *blastp*) was performed using known marker transcript peptide sequences as queries to search on the self-established sorghum protein database. Critical parameters for *blastp* are “*blastp −query known-marker-sequence −db sorghum-protein-database −max_target_seqs 5-max_hsps 1-evalue 1e-3-outfmt ’7 qseqid sseqid length pident ppos qlen slen qstart qend sstart send evalue*’”. For each marker transcript, only the best sorghum transcript hit(s), which has the lowest E-value and passes E-value threshold of 1e-3, was treated as the best hit. BLAST was performed on the isoform peptide sequence, but this result at the transcript level was finally summarized at the gene level (Supplemental Data Set 2).

### Gene Ontology (GO) enrichment analysis

We used *enricher* function in *ClusterProfiler* (G. Yu et al., 2012) package to perform GO term enrichment analysis. Prior to the enrichment analysis, a GO annotation file for sorghum was created, including not only direct GO terms but also parental GO terms (i.e., nodes at higher hierarchies) for each gene. Direct GO term annotation for sorghum was achieved from *PlantRegMap* (http://plantregmap.gao-lab.org/download.php); parental GO term annotation was achieved using *buildGOmap* function in *CluseterProfiler*; information of “Term” (description of GO term) and “Ontology” (BP, CC, MF) was retrieved from the *GOTERM* dataset in *GO.db* package (Carlson et al., 2019). Only GO terms with FDR < 0.05 in the enrichment test were thought of as enriched terms for the testing gene set. In our analysis, enrichment analysis for cell-type specific genes usually uncovered a large number of enriched GO terms (90 on average). To facilitate interpretation and visualization, these enriched GO terms were clustered if gene overlap between two individual GO terms, defined as the Jaccard similarity coefficient (Popescu et al., 2006), was more than 40%. The function of each enriched GO cluster was manually summarized. Both enriched GO clusters and orphan GO terms that cannot be connected to other GO terms were visualized in Figure 4.

### Metabolic pathway and transporter (KEGG) enrichment analysis

KEGG annotations (i.e. K number) of sorghum genes were retrieved from the *Phytozome* sorghum annotation file v3.1.1, where 8744 of 34129 genes in the genome have been annotated with K numbers. To get a more comprehensive KEGG annotation, *BlastKOALA* (Kanehisa et al., 2016) was used to assign the K number to genes that are absent of KEGG annotation. This process increased the number of sorghum genes with KEGG annotations to 10917. K numbers involved in the metabolic pathway and transporter category were achieved from KEGG database. Fisher’s exact test was implemented for enrichment analysis to estimate whether a metabolic pathway or a transporter category is enriched in cell-type specific genes, compared to genome background.

### Enriched motif analysis

Enriched motif analysis, including known and *de novo* motifs, was performed on putative promoter regions of 1500 bp upstream of ATG at the gene level. Promoter sequences of sorghum genes were retrieved using *Phytozome Biomart* tool. For known motifs, we used *PlantPan 3.0* promoter analysis tool to scan promoter regions to get all known motifs. Fisher’s exact test was performed to estimate whether genes containing a certain motif are enriched in cell-type specific genes compared to the genome background (FDR < 0.05). For *de novo* motifs, we used *MEME* to discover enriched motifs in promoter regions of cell-type specific genes. Critical parameters for *MEME* are “*meme-mod anr-nmotifs 20-minw 5-maxw 30-objfun classic-revcomp-bfile 0-order background markov model*”. A background Markov model for 1500 bp promoters of all sorghum genes was built using *RSAT create-background-tool* (http://rsat.sb-roscoff.fr/create-background-model_form.cgi). For each *de novo* enriched motif, *Tomtom* (Gupta et al., 2007) was used to find the top known motif that is most and significantly similar to it so that possible binding TFs can be estimated. Critical parameters for *Tomtom* are “*tomtom-no-ssc-min-overlap 5-mi 1-dist pearson-internal-evalue-thresh 10-incomplete-scores CIS-BP_2.00/Sorghum_bicolor.meme*”. TF families that can bind either known or *de novo* enriched motifs were summarized as a heatmap (Figure 7B), where color darkness represents the proportion of cell-type specific genes that can be bound by a TF family through enriched motif(s).

### Gene regulatory network (GRN) construction

In this study, we constructed different GRNs by integrating the co-expression strength among gene pairs and putative TF-motif interaction to investigate regulatory landscapes of a cell type and/or an interested pathways. Detail information for different GRNs on node identities, edge filtering criteria, and input dataset for calculating the co-expression strength can be found in Supplemental Data Set 12. Generally, we applied gene filtering and normalization processes for input datasets, same with that in Tau calculation (see above). Pearson Correlation Coefficient (PCC) was calculated to estimate the co-expression strength. Putative TF-motif interactions in the promoter region were achieved using *PlantPan 3.0* promoter analysis tool. Edges in GRNs, which connect source nodes and target nodes, were kept if they passed certain criteria set for PCC (Supplemental Data Set 12) and contain putative TF-motif interactions as well. We used *Cytoscape* (https://cytoscape.org/) for GRN visualization and network analysis.

### Accession Numbers

Sequence data from this article can be found in the GenBank data libraries under accession number GSE218642.

## Supplemental Data files

**Supplemental Figure 1:** Stem bulk tissue combining multiple cell types demonstrated a dilution effect for the expression of cell-type specific genes compared to single cell type.

**Supplemental Figure 2:** Distinct TF families play a dominant role in regulating common up-regulated differentially expressed genes (DEGs) in vascular bundle and non-vascular bundle cells.

**Supplemental Figure 3:** Different stem cell types of Wray were collected using LCM. **Supplemental Data Set 1:** Cell-type specific genes (0.8 ≤ Tau ≤ 1 & Wilcoxon p-value < 0.05) identified in this study with gene annotations of their Arabidopsis and rice homologs.

**Supplemental Data Set 2:** Known cell-type marker genes from previous studies associated with sorghum stem cell-type specific genes identified in this study.

**Supplemental Data Set 3:** Differentially expressed genes (DEGs) from pairwise comparisons of vascular bundle and non-vascular bundle cells.

**Supplemental Data Set 4:** Enriched Gene Ontology (GO) Biological Processes (BPs) for common differentially expressed genes (DEGs) between vascular bundle and non-vascular bundle cells.

**Supplemental Data Set 5:** Enriched Gene Ontology (GO) Biological Processes (BPs) for cell-type specific genes.

**Supplemental Data Set 6:** Cell-type specific genes involved in KEGG metabolic pathways. **Supplemental Data Set 7:** Cell-type specific genes involved in KEGG transporter categories. **Supplemental Data Set 8:** Hierarchy for TF family binding the largest proportion of cell-type specific genes through enriched motifs.

**Supplemental Data Set 9:** Expressions of *CESA4/7/8* and their co-expressed genes in the stem developmental dataset.

**Supplemental Data Set 10:** Expressions of genes involved in SCW formation in the stem developmental dataset and LCM-derived stem cell-type dataset.

**Supplemental Data Set 11:** Regulatory interactions among TFs involved in SCW formation. **Supplemental Data Set 12:** Node, edge and dataset for the construction of different gene regulatory networks (GRNs).

**Supplemental Data Set 13:** Vascular bundle and non-vascular bundle GRNs.

**Supplemental Data Set 14:** Cell-type specific GRNs.

**Supplemental Data Set 15:** Cell-type specific gene and all TFs GRNs.

**Supplemental Data Set 16:** SCW formation GRN.

**Supplemental Data Set 17:** SCW TF GRN.

## Supporting information

Supplemental Data Set

## Acknowledgements

Ka Man Jasmine Yu for collecting stem samples. Austin Lamb for performing stem cross-section FASGA staining. Ryan Tillman for assistance with LCM.

## Author Contributions

GO, JM, KS, and AMC conceived the experiments; JF, BM and HM analyzed the data; BJ, WC and MG developed LCM protocol and collected the samples; LMM prepared libraries and performed RNA-sequencing; KMJY collected plant samples; JF, BM, BJ, WC, LMM, JM, KS, and AMC wrote the manuscript; GO, JM, KS, and AMC obtained funding for the project.

Jie Fu

Brian McKinley

Brandon James

William Chrisler

Lye Meng Markillie

Matthew J Gaffrey

Hugh D Mitchell

Galya Orr

Kankshita Swaminathan

John Mullet

Amy Marshall-Colon

