## supplemental figures for "The stem cell-type transcriptome of bioenergy sorghum reveals the spatial regulation of secondary cell wall networks"

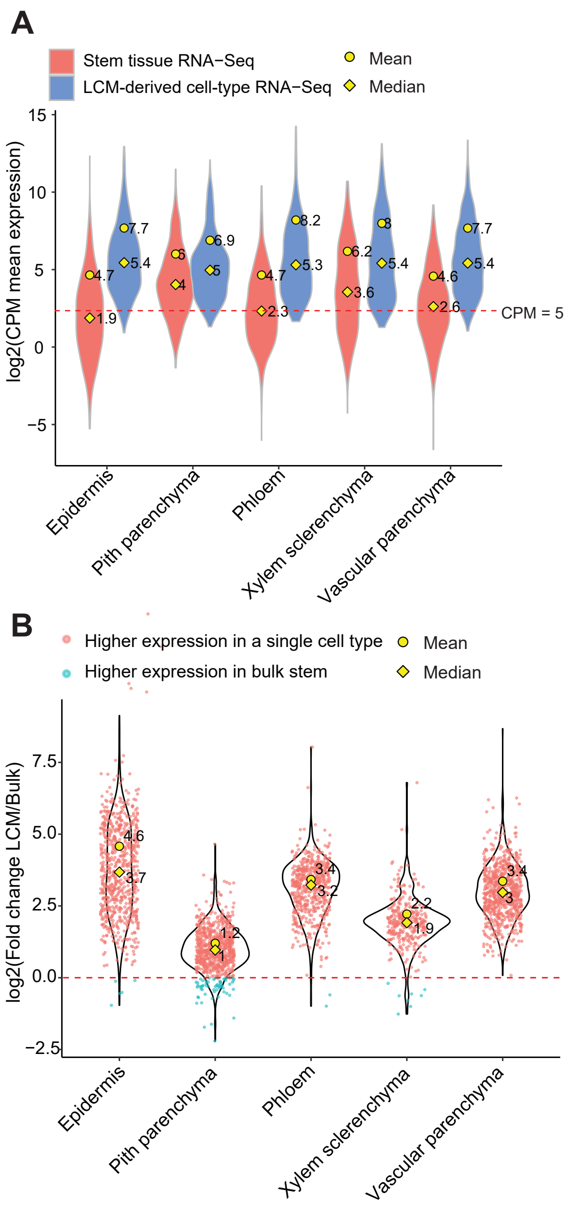


Supplemental Figure 1 Stem bulk tissue combining multiple cell types demonstrated a dilution effect for cell-type specific genes (0.8 ≤ Tau < 1 & Wilcoxon p-value < 0.05) compared to single cell type (Supports Figure 2). (A) Comparison of cell-type specific gene expressions between the stem bulk RNA-Seq (red) and LCM-derived cell-type RNA-Seq (blue). (B) Fold change of cell-type specific gene expressions between two datasets (Fold change = Expression in a single cell type / Expression in stem bulk tissue).

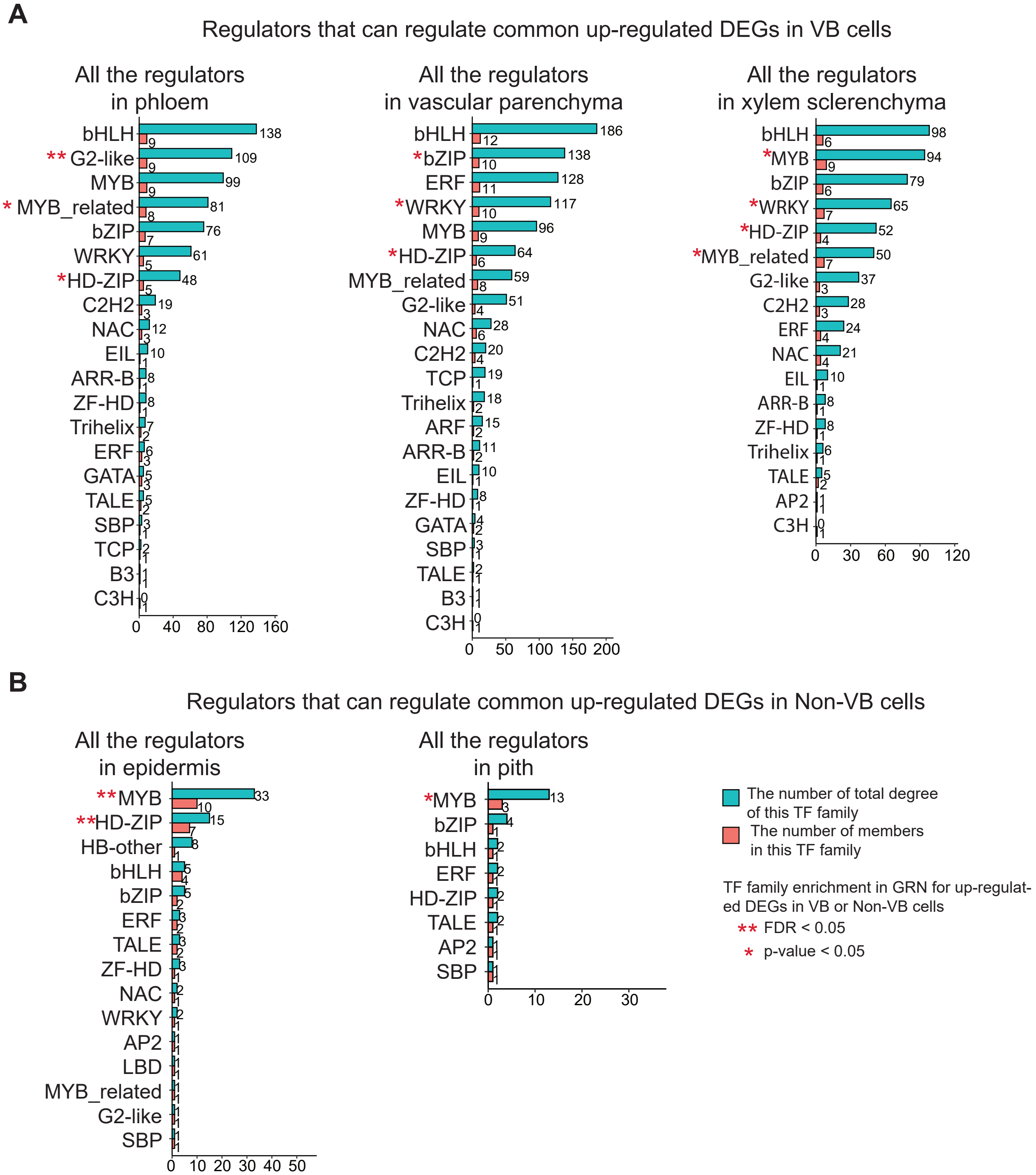


Supplemental Figure 2 Distinct TF families play a dominant role in regulating common up-regulated DEGs in vascular bundle (VB) cells (A) and non-VB cells (B) (Supports Figure 3). Estimation of enrichment for a TF family in GRN is conducted by Fisher's exact test. The GRN for a certain cell type uses the common up-regulated DEGs as target nodes (genes being regulated) and the union of up-regulated DEGs in this cell type as source nodes (regulators). For example, in phloem GRN, target nodes are 127 common differentially expressed genes in vascular bundle cells (intersection of pairwise comparisons); source nodes are up-regulated DEGs in phloem compared to either epidermis or pith parenchyma cells (union of pairwise comparisons).

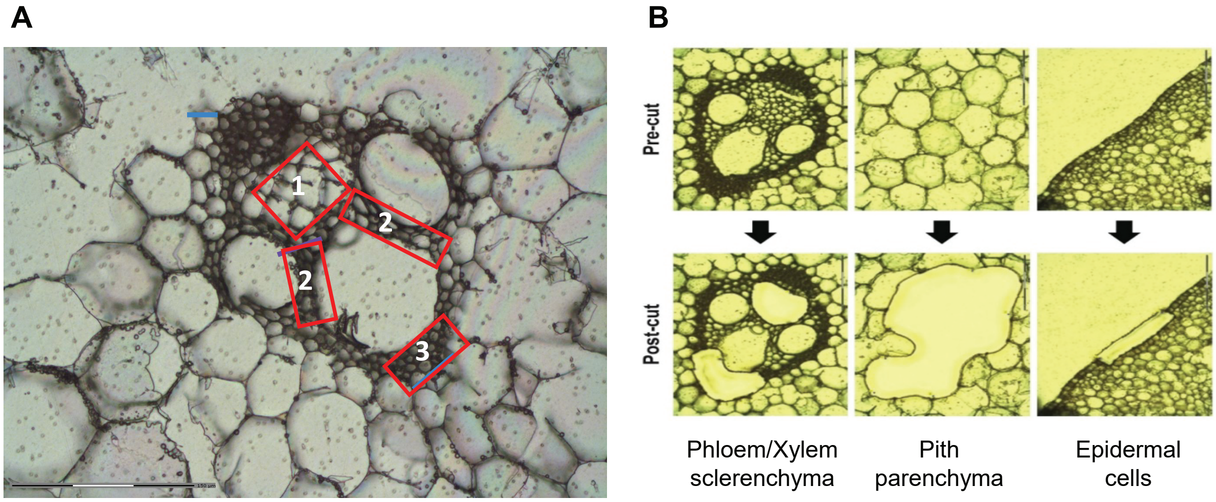


Supplemental Figure 3: Different stem cell types of Wray were collected using LCM (Supports Figure 1 and 2). (A) The anatomy of a vascular bundle, marked with positions of phloem (#1), vascular parenchyma (#2), and xylem sclerenchyma (#3). (B) Pre-cut (top panel) and post-cut (bottom panel) of stem cell types using LCM.
